# Bright calcium-modulated bioluminescent indicators for activity imaging and red photon-assisted synaptic transmission

**DOI:** 10.64898/2026.01.20.700092

**Authors:** Yufeng Zhao, Montserrat Porta-de-la-Riva, Sungmoo Lee, Yan Wu, Mark Klein, Mary P. Hall, Yichi Su, Lance P. Encell, Thomas A. Kirkland, Michael Krieg, Michael Z. Lin

## Abstract

Bioluminescent calcium sensors have unique uses in neuroscience and neuroengineering, enabling noninvasive imaging of neuronal activity and contactless activation of opsin-expressing neurons. However, the speed, range, and robustness of non-invasive imaging and rewiring or neuronal activity are limited by the radiance and dynamic range of existing bioluminescent calcium sensors. Here, we report the stepwise engineering of improved cyan-excitable red fluorescent proteins (mCyRFP4 and dCyRFP4), an improved red bioluminescent protein based on NanoLuc and CyRFP4 (Antares3), and an improved bioluminescent calcium sensor based on Antares3 (CaMBI3). Antares3 is 3-fold brighter than its predecessor, while CaMBI3 responds with an overall dynamic range of 24-fold, enabling high-sensitivity detection of calcium dynamics in muscle and neuronal tissues *in vivo*. Finally, a CaMBI3 variant and a red-shifted opsin ChRmine enabled red photon-assisted synaptic transmission in *C. elegans*. CaMBI3 thus facilitates genetically targeted non-invasive imaging and rewiring of neural activity in living animals.

## Introduction

Bioluminescence, luciferase-catalyzed light emission from a chemical substrate, has reemerged as an increasingly powerful modality for biosensor development and application. By eschewing excitation light, bioluminescence avoids multiple complications that occur with fluorescence imaging, including tissue damage from optical elements, autofluorescence, phototoxicity, and thermotoxicity. While insect luciferases paired with D-luciferin or their derivatives have been extensively used *in vivo* for basic cell tracking^1^, NanoLuc derived from a deep shrimp luciferase has proven to be a catalytic core for a wide array of bioluminescent activity indicators, including CaMBI^2^ and GeNL variants for calcium^3^ and KiMBI for kinases^4,5^. The development of substrates optimized for body and brain imaging such as fluorofurimazine (FFz)^6^ and cephalofurimazine (CFz)^7,8^ has enabled the use of these reporters to track event in animal models noninvasively. Finally, a NanoLuc-based calcium indicator was used for photon-assisted synaptic transmission (PhAST)^9^ in which calcium-induced bioluminescence in transmitter neurons activated channelrhodopsins in receiver neurons to create contact-free neuronal transmission.

However, bioluminescent sensors are limited in their applications by low output intensity and dynamic range. While NanoLuc exhibits the highest known photon production rate of bioluminescent systems characterized so far, it exhibits a lower rate of photons produced per enzyme molecule than fluorescent proteins do (hundreds per second compared to millions). To preserve bioluminescence signal from deep locations in the body, longer emission wavelengths are preferable for their reduced scattering and absorption by biological tissues. Previously, we developed the bright cyan-excitable orange-red fluorescent proteins CyOFP and CyRFP to red-shift NanoLuc emission via Förster resonance energy transfer (RET, also known as bioluminescence resonance energy transfer or BRET)^10,11^. Antares1, a fusion of two CyOFP1 domains with NanoLuc, and Antares2, in which NanoLuc is replaced by its derivative TeLuc, emit a substantial fraction their photons at orange and red wavelengths, enabling sensitive deep-tissue imaging in mammalian subjects^10,12^. However, the emission spectra of Antares1 and Antares2 still peak below 600 nm. Further improvements of Antares for emission intensity and red-shifting would be desirable to enhance the speed, sensitivity, or robustness of cell tracking, calcium reporting, and related methods such as PhAST. Here, we present a suite of genetically encoded fluorescent and bioluminescent tools with enhanced performance for a broad range of imaging applications. First, we engineered mCyRFP4 and dCyRFP4 which feature 20-nm red-shifted emission peaks relative to CyOFP and CyRFP3. Second, building upon dCyRFP4, we developed an improved bioluminescent reporter, Antares3, which displays a threefold increase in cellular brightness and a fourfold enhancement in red photon emission compared to Antares1. Third, we inserted calcium-sensing domains into Antares3 and performed directed evolution to obtain CaMBI3 as an improved calcium indicator. We comprehensively characterized CaMBI3 *in vitro* and *in vivo*, demonstrating its superior brightness and dynamic range for Ca^2+^ reporting than CaMBI. Finally, we used CaMBI3 and the red-absorbing opsin ChRmine to establish the first red-shifted PhAST system, enabling successful neurotransmission using red photons.

## Results

### Engineering CyRFP variants with red-shifted emission

To boost the amount of red photons (>600 nm) emitted by Antares1, we first engineered a red-shifted cyan-excitable red fluorescent protein. In Antares1, two CyOFP domains, whose emission peak is at 586 nm, are used as acceptors for NanoLuc^10^. Further efforts to improve CyOFPs led to red shifted variants named mCyRFP1^13^ and mCyRFP3^11^, but neither exhibits emission peak above 600 nm (594 nm for mCyRFP1 and 588 nm for mCyRFP3).

We used structure-guided and random mutagenesis combined with bacterial colony screening to further red-shift mCyRFP3 emission (**Supplementary Fig. 1a**). Randomization of H28, a key position in red-shifting in mCardinal^14^, yielded H28A with a red-shifted emission peak at 598 nm (**Supplementary Fig. 1b**,**c**). Randomizing T60 and M61 simultaneously, we discovered a H28A T60Q M61S mutant with a 20-nm red-shifted emission peak of 608 nm but decreased brightness (**Supplementary Fig. 1b**,**c**). A further round of random mutagenesis yielded mCyRFP3 H28A T60Q M61S R122I H169Y (**Supplementary Fig. 1d**) with restored brightness. This mutant contains a hydrophobic mutation at R122I, which resides in a dimeric interface in the ancestral coral protein, so was expected to be dimeric at higher concentrations. Random mutagenesis of this protein in the context of Antares (described below) yielded G49D A104V with further improved brightness. This variant was indeed dimeric at 10 μM **(Supplementary Fig. 1e)**, and was designated dCyRFP4. Compared to CyOFP, dCyRFP4 exhibits a similar excitation spectrum but its emission spectrum is red-shifted by 22 nm (**Fig. 1a, Table 1**).

**Table 1.** Characteristics of purified CyOFP and its variants.

| Mutant | Extinction coefficient ($M^{-1} cm^{-1}$) | Quantum yield | Emission peak (nm) | Oligomeric state at 10 $\mu M$ |
| --- | --- | --- | --- | --- |
| CyOFP | 40,000 | 0.76 | 586 | Dimer |
| mCyRFP3 | 38,000 | 0.72 | 588 | Monomer |
| dCyRFP4 | 28,000 | 0.71 | 608 | Dimer |
| mCyRFP4 | 31,000 | 0.55 | 608 | Monomer |

**Figure 1.**
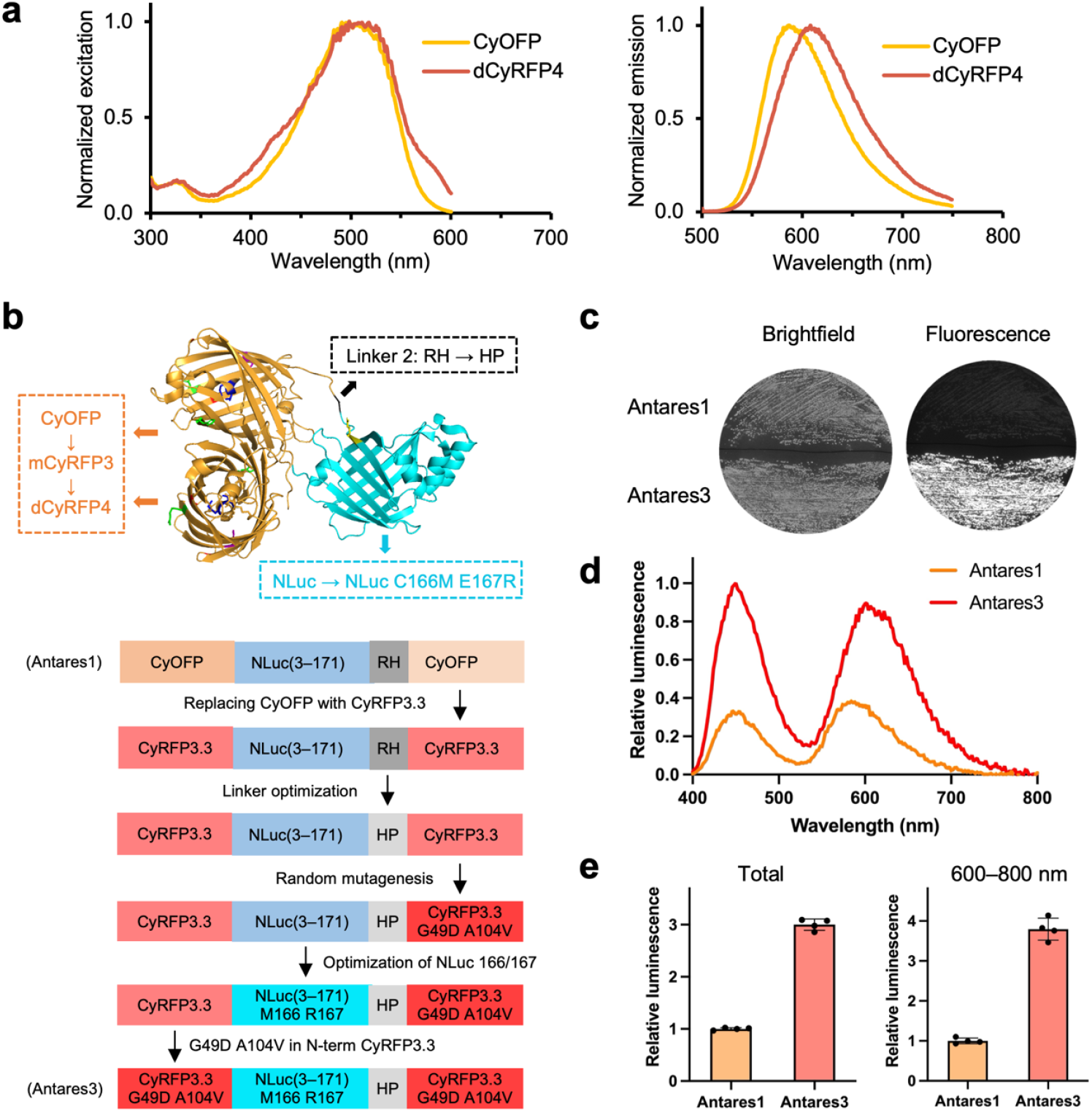
Development and characterization of Antares3. **a**, Normalized excitation and emission spectra of purified CyOFP and dCyRFP4. **b**, Structural model of Antares3 (top) and engineering history (bottom). The model is based on crystal structures of NanoLuc (cyan, PDB 5IBO) and CyOFP (orange, 5BQL). Mutations of Antares3 relative to Antares1 are shown in other colors. **c**, Brightfield and fluorescence images of patches of bacteria expressing Antares3 and Antares1. **d**, Relative luminescence spectra of Antares3 and Antares1 using FFz measured in the diluted lysate of HEK293A cells transfected with the same amount of DNA. **e**, Total luminescence intensity (left) or red emission intensity (right) was measured with readings at 10 timepoints spaced 1 minute apart, begun immediately after FFz addition. This time interval was chosen to contain nearly the entire FFz conversion time course. Error bars represent standard deviation.

To generate a more strongly monomeric cyan-excitable fluorescent protein with the desirable bathochromic emission of dCyRFP4, we selected additional bright mutants from random mutagenesis of mCyRFP3 H28A T60Q M61S, discovering a K25M V95M A104V Y211F variant (**Supplementary Fig. 1a**,**d**), that was monomeric on size-exclusion chromatography at 100 μM **(Supplementary Fig. 1e)**, while maintaining an emission peak at 608 nm (**Table 1, Supplementary Fig. 2a**). We designated this protein mCyRFP4.

### Ratiometric voltage imaging with mCyRFP4

The increased Stokes shift of mCyRFP4 should be advantageous for single-excitation dual-emission imaging with green fluorescent reporters. We had previously fused the green fluorescent genetically encoded voltage indicator (GEVI) ASAP3^15^ to mCyRFP3 so the voltage-independent red fluorescence of mCyRFP3 could serve as a reference channel for motion correction^11^. We subsequently created ASAP4e as a GEVI that increases brightness with positive changes in membrane potential, the opposite of ASAP3, resulting in improved photostability^16^. A ratiometric positively tuned GEVI based on ASAP4e and mCyRFP3 would be desirable, but ASAP4e exhibits red-shifted emission relative to ASAP3, increasing overlap with mCyRFP3. As the redder emission of mCyRFP4 should be more easily separated from ASAP4e emission (**Supplementary Fig. 2a**), we hypothesized mCyRFP4 would allow larger changes in green/red fluorescence ratios with depolarization. To test this, we created fusions of ASAP4e at its C-terminus with mCyRFP3 and mCyRFP4, resulting in ASAP4e-R3 and ASAP4e-R4 (**Supplementary Fig. 2b**). When imaged using a dual-emission optical setup at a single excitation wavelength (**Supplementary Fig. 2c**), ASAP4e-R4 achieved a 75% higher green-channel intensity response to a voltage step from –70 mV to +30 mV in patch-clamped HEK293A cells compared to ASAP4e-R3 (mean ΔF/F of 159% vs. 91%, **Supplementary Fig. 2d**), as expected from reduced contaminatino of mCyRFP4 emission into the green channel. Most importantly, the green-to-red ratiometric response for ASAP4e-R4 was substantially higher than ASAP4e-R3 (mean ΔR/R of 143% vs. 85%, **Supplementary Fig. 2e**).

### Engineering brighter red bioluminescent proteins

To create a red-shifted Antares, we carried out a systematic mutagenesis and screening strategy involving optimization of all three domains in the protein (**Fig. 1b**). First, we replaced either or both of the two CyOFP domains of Antares1 with mCyRFP3 H28A T60Q M61S R122I H169Y (CyRFP3.3) and assessed RET and bioluminescent brightness in mammalian cell lysates (**Supplementary Fig. 3a**). CyRFP3.3-NanoLuc-CyRFP3.3 produced the brightest red emission (>70% more photons above 600 nm than Antares1), despite lower RET efficiency compared to Antares1 (**Supplementary Fig. 3b**), and was selected for further evolution with substrate FFz. Rescreening the linker at the C-terminus of NanoLuc, we found linker sequence His-Pro to slightly improve RET **(Supplementary Fig. 3c)**. Screening random mutants throughout the protein for further RET improvement then yielded mutations G49D and A104V in the C-terminal CyRFP domain to create the above-described dCyRFP4 **(Supplementary Fig. 3c)**. Finally, overall brightness was improved by mutations M166 and R167 in the NanoLuc domain, obtained by re-screening of C166X E167X, and by converting the N-terminal CyRFP domain to dCyRFP4 **(Supplementary Fig. 3d)**. This variant, designated Antares3, was notably brighter compared to Antares1 in bacteria (**Fig. 1c**). It also exhibits a 22-nm bathochromic shift in its red emission peak (**Fig. 1d**).

We compared the performance of Antares3 to Antares 1 in multiple mammalian cell lines after transient transfection and FFz addition. Antares3 was brighter than Antares1 by a factor of ~3 (all wavelengths) or ~4 (> 600 nm) in HEK293A cell lysates (**Fig. 1e**). Antares3 was also brighter than Antares1 in HeLa and Huh-7 cells (**Supplementary Fig. 4**). The combined improvement in brightness and red-shifted emission are advantageous features for deep imaging through mammalian tissue.

To explore alternative schemes for red-shifting NanoLuc, we tested a RET relay approach by fusing GeNL^3^, the brightest green bioluminescent protein, to mScarlet-I^17^ (**Supplementary Fig. 5a-c**). GeNL uses mNeonGreen as acceptor for NanoLuc and it is 80% brighter than NanoLuc due to the high quantum yield of mNeonGreen. mScarlet-I is known to be a best FRET acceptor of mNeonGreen as demonstrated by others^18^. We fused it to the N-term of GeNL as an acceptor to increase the number of emitted red photons. A proline was used as a linker which allows the domains on its both sides to fold independently. Indeed, Scarlet-P-GeNL generates 50% (whole spectrum) and 64% (> 600 nm) more photons than Antares1 when expressed in HEK293A cells (**Supplementary Fig. 5d**). We screened the positions C166X E167X of the NanoLuc domain against substrate FFz and found that Scarlet-P-GeNL C166M is 2.5-fold (whole spectrum) and 2.7-fold (> 600 nm) of Antares1. While Scarlet-P-GeNL C166M is brighter than Antares1, it is dimmer compared with Antares3 **(Supplementary Fig. 5d-f**).

### Engineering a high performance Antares3-based calcium indicator

We asked whether the beneficial mutations from the evolution can be generalized to the Antares1-derived calcium indicator CaMBI-110 (hereafter referred to as CaMBI1, **Fig. 2a**). Therefore, we replaced Antares1 with Antares3 as the scaffold of CaMBI1 and compared the new indicator with the original one in cultured cells. Given the widespread application of Ca^2+^ imaging in the CNS, we chose the CNS-optimal substrate CFz for the initial evaluation and optimization of the new CaMBI sensor. As expected, we found that Antares3-CaMBI exhibited 2-fold improved signals, and did not compromise its ability to sense calcium (**Supplementary Fig. 6**). Thus, Antares3 is a generally applicable brighter substitute for Antares1, and the new CaMBI variant is designated CaMBI2.5.

**Figure 2.**
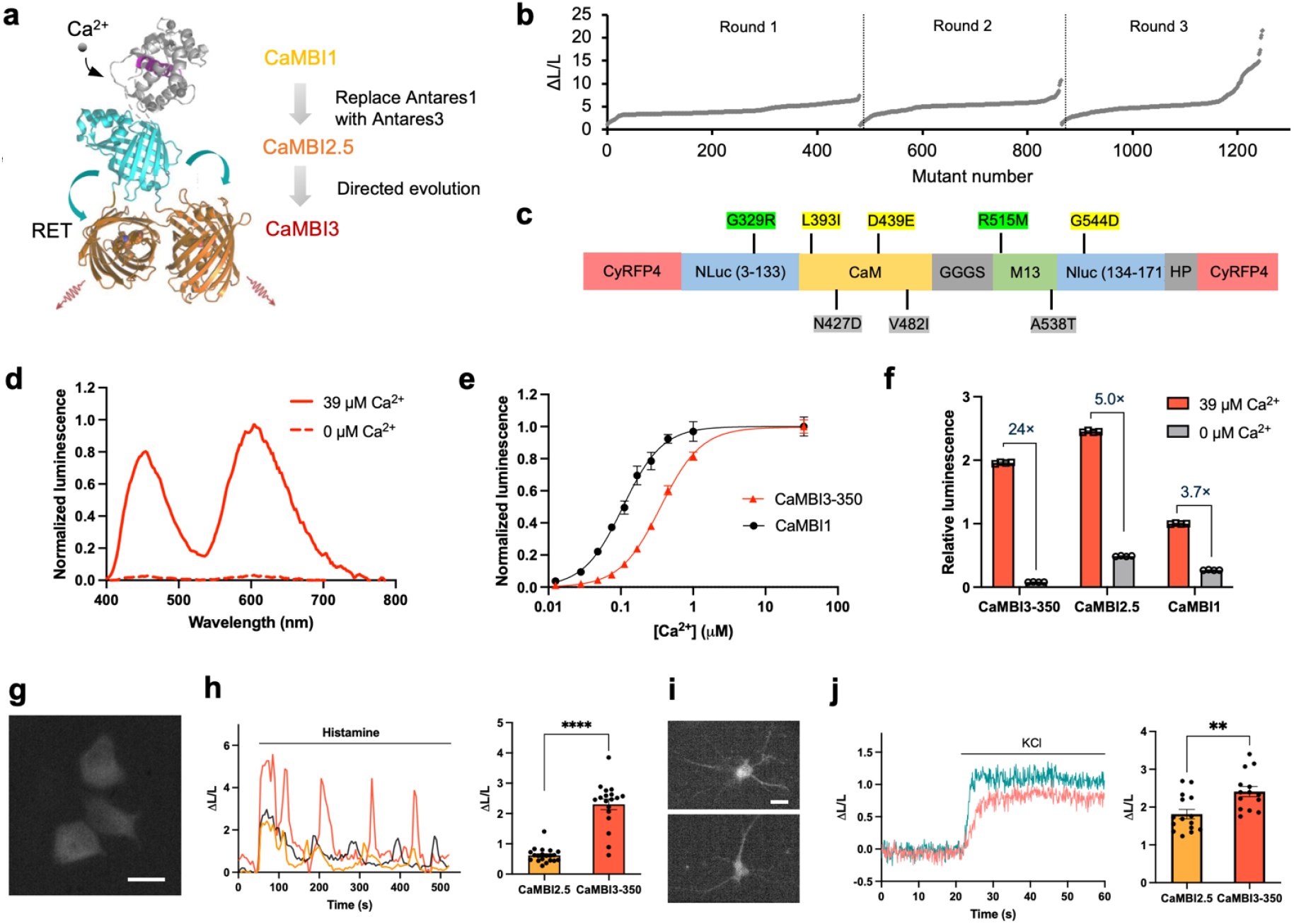
Development and characterization of CaMBI3. **a**, Schematic structure and evolution of CaMBI3 from CaMBI1. **b**, Summary of responsiveness of CaMBI variants screened in three rounds of directed evolution. **c**, Mutations of CaMBI3 relative to CaMBI1. Yellow highlights: 1^st^ round. Green highlights: 2^nd^ round. Grey highlights; 3^rd^ round. **d**, Luminescence spectra of CaMBI3 in diluted bacteria lysates in the presence or absence of calcium.. **e**, Titration of CaMBI3 in diluted bacterial lysates with calcium standard solutions. **f**, Comparison of the responsiveness of CaMBI1, CaMBI2.5 and CaMBI3 in mammalian cell lysates. Error bars represent standard deviation. **g**, Luminescence image of HeLa cells expressing CaMBI3. **h**, Left: representative luminescence traces of CaMBI3 in response to histamine induced calcium oscillation. Right: comparing CaMBI2.5 and CaMBI3 for the dominant signal increase upon histamine addition. Error bars represent s.e.m. **i**, Luminescence image of cultured neurons expressing CaMBI3. **j**, Representative luminescence traces (left) and comparison of CaMBI2.5 and CaMBI3 in dissociated neurons (right) in response to KCl induced depolarization. Error bars represent s.e.m.. Scale bars in (**g**) and (**i**):20 µm. CFz substrate was used for all measurements.

We next sought to improve the dynamic range of CaMBI2.5. We reasoned that mutations in the NanoLuc and calcium-sensing moieties could alter the energetics of calcium-induced conformational changes to improve dynamic range. We performed error-prone PCR to these regions, screened 400–500 randomly selected variants for luminescence in 0 and 39 μM calcium, and used the best-performing variants as templates for additional rounds of mutagenesis and selection. After three rounds of directed evolution (**Fig. 2b**), we obtained an improved variant with two mutations in NanoLuc and six in CaM-M13 (**Fig. 2c**, and **Supplementary Fig. 7**). This variant demonstrated 24-fold dynamic range between calcium-free and -saturated states in diluted cell lysates (**Fig. 2d**) and a *K*_d_ of 230 nM with FFz and 350 nM with CFz, compared to 85 nM (FFz) and 110 nM (CFz) for CaMBI1 (**Fig. 2e**). We thus named this variant CaMBI3-350.

CaMBI3-350 also exhibited approximately 5-fold enhanced dynamic range compared to the original template CaMBI2.5 in mammalian cell lysates (**Fig. 2f**). The cellular brightness of CaMBI3-350 in the Ca^2+^-bound state (39 μM Ca^2+^) was 96% higher than that of CaMBI1, suggesting either greater intracellular abundance or higher per-molecule photon production. Although CaMBI3 was evolved using CFz as the substrate, we tested both CFz and FFz in mammalian cell lysates and observed no significant difference in performance (**Table 2**).

**Table 2.** Summary of in vitro properties of CaMBI variants.

| Mutant | $K_d$ (CFz-FFz, nM) | Dynamic range in HeLa lysates (FFz) | Dynamic range in HeLa lysates (CFz) | Histamine induced response (N) | KCl depolarization neuron (N) |
| --- | --- | --- | --- | --- | --- |
| CaMBI1 | 110–85 | 3.8 | 3.7 | 0.59±0.38 (17) | ND<sup>1</sup> |
| CaMBI2.5 | ND<sup>1</sup> | 4.2 | 5 | 1.36±0.34 (15) | 1.77±0.39 (15) |
| CaMBI3-350 | 350–230 | 22 | 24 | 2.69±0.89 (12) | 2.4±0.5 (14) |
| CaMBI3 H362Y M515R | 280–200 | 26 | 30 | 2.37±0.69 (14) | 2.16±0.64 (16) |
| CaMBI3 H362Y I393L M515R | 260–150 | 24 | 27 | 1.71±0.59 (13) | 1.99±0.47 (14) |

<sup>1</sup> Not determined.

To eliminate unnecessary mutations in the course of directed evolution, we performed single mutation reversion of all the eight mutations and measured the dynamic range of each variant (**Supplementary Fig. 8a**). We also introduced a beneficial mutation, H362Y, discovered during directed evolution that did not exist in CaMBI3-350. As a result, we obtained two new variants, CaMBI3 H362Y M515R and CaMBI3 H362Y I393L M515R, with small increases in dynamic range and calcium affinity (**Table 2**), and no change in bioluminescent resonance energy transfer efficiency (**Supplementary Fig. 8b**).

We also investigated whether the improved dynamic range of the three CaMBI3 variants is retained in the absence of the CyRFP4 BRET acceptors. Blue CaMBI3 (bCaMBI3) versions lacking any fluorescent protein fusion were constructed from each of the CaMBI3 variants. High signal contrast was observed for all three variants when bacterial lysates containing the proteins were diluted into calcium buffer with CFz substrates (**Supplementary Fig. 9**). These results confirm that removal of the CyRFP4 domains does not affect the dynamic range of CaMBI3.

We next compared the responses of CaMBI2.5 and CaMBI3 variants to cytosolic calcium influx in cultured mammalian cells. We first expressed CaMBI variants in HeLa cells through transient transfection (**Fig. 2g**) and recorded luminescence responses upon histamine stimulation. The first calcium wave was used to evaluate the performance of the indicators. CaMBI3-350 demonstrated higher responses than CaMBI2.5 (**Fig. 2h**). To assess the sensitivity of the CaMBI variants in neurons, we expressed the variants in cultured dissociated rat neurons (**Fig. 2i**) and recorded their response to elevated calcium levels upon KCl-induced neuronal depolarization (**Fig. 2j**). Among all the variants examined, CaMBI3-350 demonstrated the largest responses to histamine or KCl stimuli in HeLa cells or dissociated neurons than other variants (**Table 2**).

### CaMBI3-350 reports calcium dynamics in muscles and neurons in *C. elegans*

To characterize the performance of CaMBI3-350 as a bioluminescent calcium reporter *in vivo*, we used *C. elegans* as an established test-bed for calcium imaging in freely behaving animals^19^. We have previously constructed a microscope capable of bioluminescent calcium imaging with sub-second exposure times. Using the transgenic line established above, we collected images of individual animals undergoing dorsoventral body swing, akin to their endogenous crawling behavior, while being mounted on a soft agar-bedding (as described previously^20^). As expected, calcium intensity closely followed body curvature (**Fig. 3a**). Similarly to before, we correlated intensity and body curvature and observed a higher bioluminescent signal on the contracted side (positive curvature) of the muscles (**Fig. 3b, c**), indicative for an increase in calcium concentration. However, we also observed substantial out-of-phase signals, suggesting that calcium activity in the muscles preceded contractile activity.

**Figure 3.**
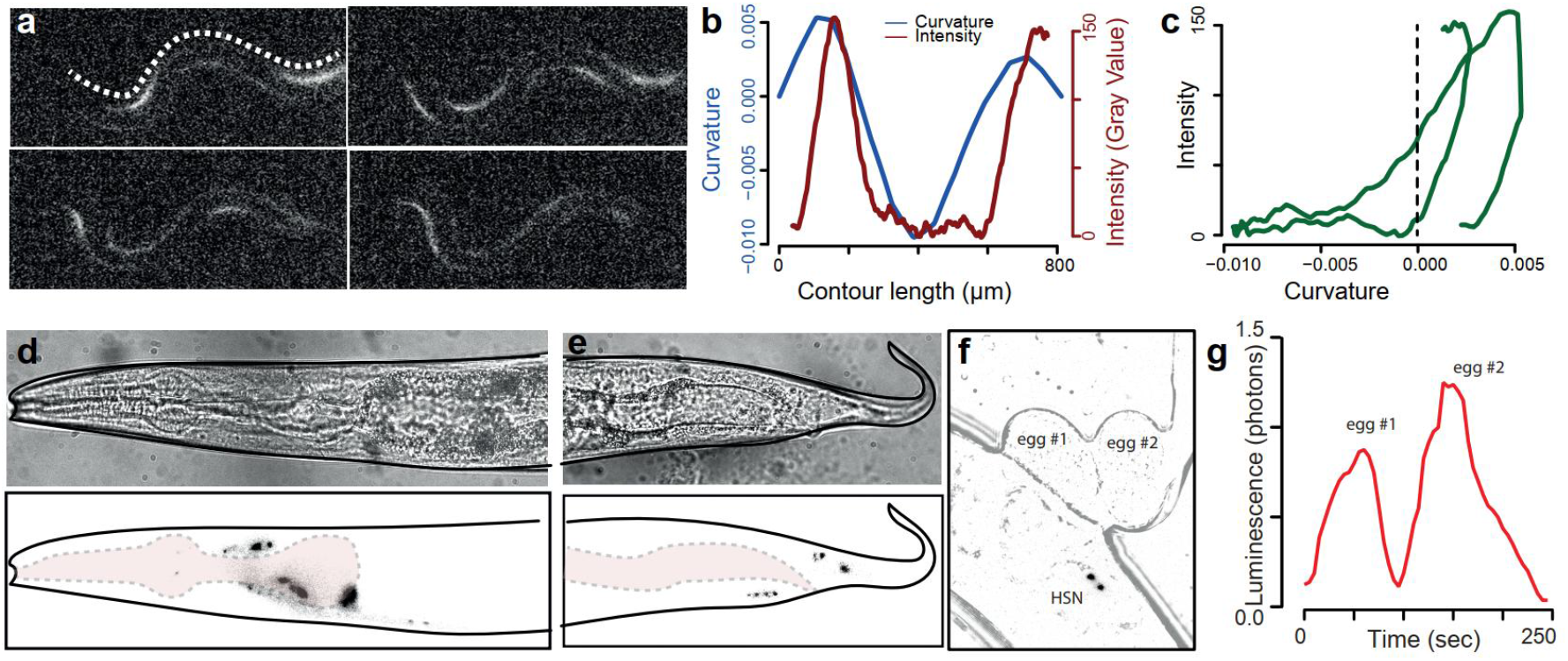
CaMBI3-350 can report calcium fluctuations in muscles and neurons. **a**, Representative still images of a moving C elegans animal expressing CaMBI3-350 in body wall muscles. The dotted line indicates the intensity profile and curvature displayed in panel (**b**). **b**, Plot of the intensity and the curvature along the length of the worm depicted by the dotted line in (**a**). **c**, Correlation of the intensity and the curvature along the length of the worm depicted by the dotted line in (**a**). **d**, Representative luminescence image of immobilized worms expressing CaMBI3-350 in head neurons (lower panel) and the corresponding brightfield image (upper panel). **e**, Representative luminescence image of immobilized worms expressing CaMBI3-350 in tail neurons (lower panel) and the corresponding brightfield image (upper panel). **f**, Representative inverted luminescence image of worms expressing CaMBI3-350 in HSN with an overlay of the corresponding brightfield image, where recently laid embryos can be observed. **g**, calcium transients of the HSN neurons in (**f**) recorded during the two egg laying events.

Next, to evaluate the performance of CaMBI3-350 as a bioluminescent reporter of neuronal activity, we generated new lines expressing CaMBI3-350 under a pan-neuronal promoter. We observed expression in many neurons in the nerve ring, mid body and the tail (**Fig. 3d, e**). Recording short movies of egg-laying neurons (**Supplementary video**), we observed a dramatic increase in HSN activity during two sequential egg-laying events (**Fig. 3f, g**). Together, these results demonstrate that CaMBI3-350 reports calcium dynamics in muscles and neurons during physiological processes in freely moving animals.

### A CaMBI3-970 variant for improved photon-assisted synaptic transmission

Low baseline activity is especially useful in bioluminescent calcium indicators, as selective activation of indicators by excitation light is not possible as it is with fluorescent indicators. When a population of cells expresses a bioluminescent calcium indicator, constitutive background photon production can transmit and scatter broadly from any expressing cell to the point of photon collection. Low background activity can thus be a highly useful feature for bioluminescent calcium indicators. This is especially true if the indicators are to be linked to activation of optogenetic effectors, such as in the method of photon-assisted synaptic transmission (PhAST).

Baseline activity of an intensiometric calcium indicator is related to its calcium affinity relative to basal concentrations. Intracellular Ca^2+^ in neurons is estimated at 100–200 nM at rest, rising to the low micromolar range upon activation.^21,22^ In previous work, we showed that Ca^2+^ affinity can be tuned by the linker length between CaM and M13, with shorter linkers reducing affinity^2^. Specifically, CaMBI1 with a GGGS linker has a *K*_d_ of 110 nM, whereas a single-glycine linker increases the *K*_d_ to 300 nM. Because CaMBI3-350 inherits the GGGS linker from CaMBI1, we hypothesized that shortening the linker would further reduce its Ca^2+^ affinity. Indeed, a CaMBI3 with a single-glycine linker exhibited a higher *K*_d_ of 970 nM (**Fig. 4a**) while maintaining a high dynamic range (**Fig. 4b**), and thus was designated CaMBI3-970. CaMBI3-970 exhibited a substantially larger luminescent response between the physiological concentraitons of 110 nM and 1.0 µM (Δ*L*/*L* = 3.9 for CaMBI3-970 vs. 1.8 for CaMBI3-350).

**Figure 4.**
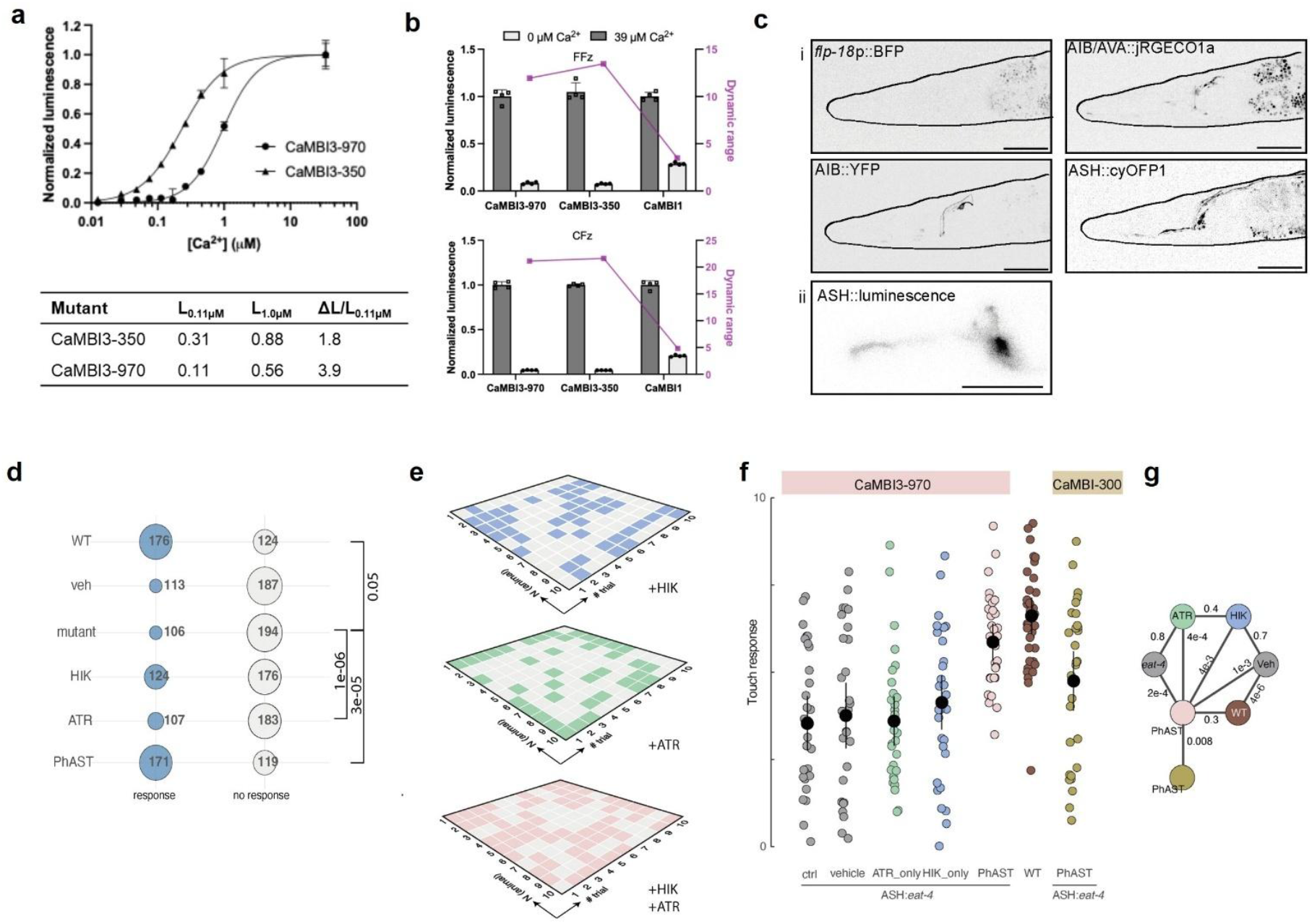
Generation of CaMBI3-970 and comparison with CaMBI1-300 in vivo. **a**, Characterization of a CaMBI3-970, a CaMBI3-350 variant with the CaM-M13 linker reduced to a single G. Top, calcium titration of CaMBI3 variants in diluted bacterial lysates with FFz. Bottom, luminescence of CaMBI3-350 and CaMBI3-970 at 0.11 and 1.0 µM in vitro, with each measurement expressed relative to the calcium-saturated luminescence. ΔL/L_0.11µM_ was calculated as (L_1.0µM_ - L_0.11µM_)/L_0.11µM_. **b**, Comparing dynamic range of CaMBI3-970, CaMBI3-350 and CaMBI1 in bacterial lysates using FFz (top) and CFz (bottom). Luminescence of each mutant was normalized to signal at 39 µM calcium. Dynamic range (purple) was calculated as L_39µM_ /L_0µM_. **c**, Fluorescent images of the PhAST components expressed in ASH, AIB and AVA (i), and 1-s luminescent image of ASH expressing CaMBI3-970. Scale bars = 50 µm. **d**, Contingency matrix for all conditions tested with CaMBI3-970. Selected p-values derived from Fisher’s exact test are indicated on the bracket to the right. **e**, Representative raster plots showing the distribution of responses and no responses for the three indicated conditions. Note, no history or adaptation is observed for individual animal touch consecutively. **f**, Scatterplot showing the average nose touch responses of each animal to 10 touches carrying a conditional eat-4 KO defect in ASH and expressing CaMBI3-970 and CaMBI1-300 in ASH. Individual datapoints (n = 30 animals) are displayed for each condition. Each data point corresponds to the average of ten touch tests per animal. Black points, medians. Vertical bars, 95% CI. **g**, Statistics for CaMBI3-970 between the indicated conditions in which the numbers indicate the two-sided P value derived from a pairwise comparison of the conditions indicated by the color code with a non-parametric Dunn test.

We previously demonstrated that cyan calcium-dependent photons from Turquoise-enhanced Nanolantern (TeNL) can be harnessed as ‘neurotransmitters’ to trans-neuronally activate light-gated ion channels. Our previous work demonstrated proof of principle with ChR2 HRDC, which demonstrates persistent cationic photocurrents after blue light absorption, and GtACR1, a blue light-responsive anionic channelrhodopsin^9^. However, red photons would have advantages in improved tissue penetration, potentially allowing for longer-range communication, and can couple to the red-absorbing opsins such as ChRmine which features high photocurrents and fast kinetics. We thus asked whether CaMBI3-970 could serve as a red-shifted alternative to TeNL, taking advantage of our previous work involving the *C. elegans* nose touch circuit. The mechanosensory neuron ASH activates interneurons AIB and AVA to mediate touch sensation (**Fig. 4c**). Wild-type *C. elegans* reverses direction when its nose is touched by an eyebrow hair, but this response is absent in a mutant deficient for glutamate, specifically in ASH (**Fig. 4d and 4e**). We created stable transgenic lines expressing CaMBI3-970 in glutamate-deficient ASH neurons and ChRmine in AIB and AVA interneurons. Intriguingly, when we reared animals in presence of both the furimazine pro-substrate hikarazine and all-trans retinol (conditions for PhAST), we observed a nearly complete rescue of the nose touch defect to levels similar to wild-type animals (**Fig. 4f**). In contrast, when tested the same assay with a transgenic animal expressing CaMBI1-300 instead of CaMBI3-970 in ASH, rescue was incomplete and not statistically significant (**Fig. 4f and 4g**; **Supplementary Fig. 10**). These results indicate that CaMBI3-970 provides superior performance over CaMBI1-300 for trans-neuronal activation of the red-absorbing opsin ChRmine.

## Discussion

In summary, using semi-rational design and directed evolution, we engineered improved cyan-excitable red fluorescent proteins (mCyRFP4 and dCyRFP4), which we then used to generate an improved red-emitting luciferase (Antares3), which we then used to create an improved red-emitting bioluminescent calcium sensor (CaMBI3). We finally combined CaMBI3 with the red-absorbing opsin ChRmine to perform red-shifted photon-assisted synaptic transmission (PhAST). Interestingly, this campaign toward improving bioluminescent sensing and rewiring of neuronal activity involved optimizing a combination with a new protein domain at each step: Starting from CyRFP4, we sequentially added NanoLuc, then calmodulin-M13, then ChRmine (**Fig. 5**). In addition, we engineered ASAP4e-mCyRFP4 as a positively tuned ratiometric GEVI. Both ASAP4e and mCyRFP4 have excitation peaks > 500 nm, and thus may allow two-photon excitation with high-power 1030-nm ytterbium-doped pulsed fiber lasers^23^. Recent advances in large-scale connectomics across multiple species have created powerful new frameworks for probing brain function at synaptic resolution^24–26^. As circuit maps grow in complexity, there is increasing demand for tools that can modulate neuronal activity with both spatial and molecular specificity. Existing neuromodulation strategies—chemical, electrical, magnetic, acoustic, and optical—offer varying degrees of resolution and invasiveness^27^. Among these, optogenetics stands out for its ability to precisely target defined neural populations, but it typically requires external light delivery through implanted devices due to the poor light sensitivity of channelrhodopsins and the strong scattering properties of brain tissue^28,29^. PhAST offers an alternative strategy wherein light is generated directly at the synapse via calcium-dependent bioluminescent luciferases, triggering light-sensitive ion channels in nearby membranes. This circuit-localized approach reduces reliance on invasive optics while retaining synapse-level precision. Previously, we demonstrated that TeNL, a high-output turquoise luciferase, can act as a “photonic neurotransmitter” to modulate nearby neurons via an inhibitory channelrhodopsin or an excitatory step-function channelrhodopsin^9^. The combination of CaMBI3-970 and ChRmine offer potential advantages. First, CaMBI3-970 background luminescence is low, allowing lower levels of baseline PhAST activity. Second, the red emission of CaMBI3-970 can penetrate deeper into mammalian tissue, raising the possibility of longer-distance neuronal relays. Third, ChRmine offers high photocurrents and rapid kinetics, allowing more sensitive responses while remaining locked temporally to the stimulus.

**Figure 5.**
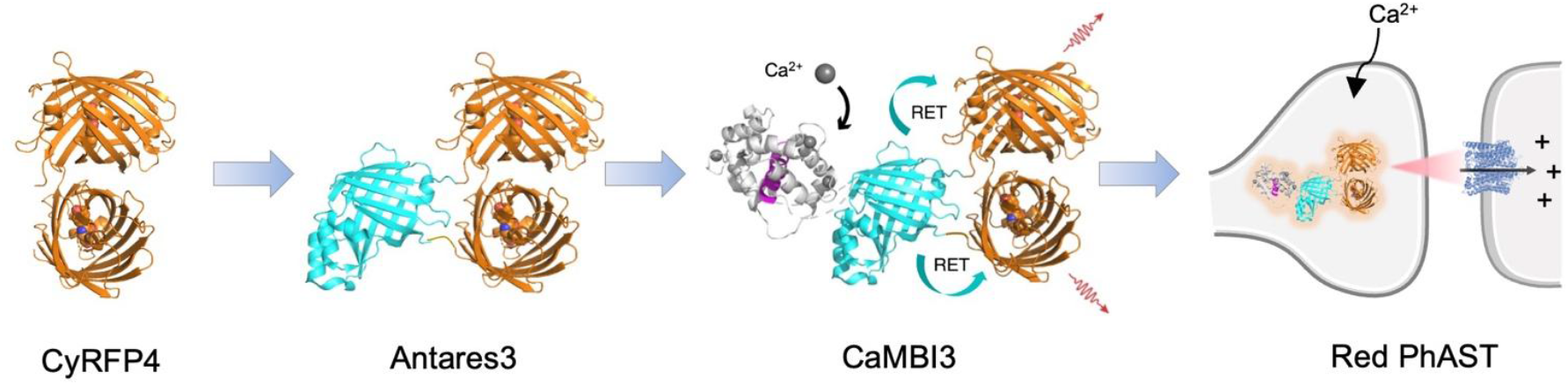
Summary of the step-wise engineering toward a high-performance bioluminescent Ca^2+^ indicator and a red PhAST system.

An examination of structures of fluorescent proteins related to mCyRFP4 yields an interesting hypothesis regarding the physical mechanism of its emission red-shift. Our first essential mutation was H28A, which red-shifted emission by 12 nm relative to mCyRFP3. In the mCyRFP3 predecessor CyOFP0.5 (PDB ID: 5BQL)^10^, H28 and the nearby Asn-41 side chains are oriented facing toward a carbonyl oxygen which acts as the terminal electron acceptor in the push-pull conjugated pi system of the chromophore. These side chains, however, are too distant to donate hydrogen bonds to the oxygen (4.1 and 4.0 Å). In our far-red fluorescent protein mNeptune, a progenitor of CyOFP, the amino acid at position 41 is glycine, and a water molecule occupies the cavity between this glycine and the carbonyl oxygen. This water molecule likely donates a hydrogen bond to the oxygen, reducing the energetic difference between ground and excited states and thereby red-shifting both excitation and emission spectra^14^. In our mNeptune-derived mCardinal, Gly-41 was replaced with Gln-41, which hydrogen-bonded with the carbonyl oxygen even more efficiently (2.5 vs. 2.8 Å), resulting in further red-shifted spectrum. We speculate that, in mCyRFP3, the mutation H28A creates a cavity that either accommodates a water molecule to hydrogen-bond with the chromophore oxygen, restoring the red-shifting mechanism of mNeptune, or allows Asn-41 to directly hydrogen-bond with the oxygen, mimicking the red-shifting mechanism of mCardinal. As mCyRFP4 demonstrates primarily an emission red-shift with minimal change in excitation spectrum, the hydrogen bond from water or Asn-41 may be induced only after chromphore excitation, similar to excited-state proton transfer in wild-type *Aequoria* GFP.

While structure-guided design is an effective strategy for tuning spectral properties, introduced mutations often decrease fluorescent protein expression or molecular brightness. Random mutagenesis is often performed to restore effective brightness, which could occur by boosting either expression or molecular brightness. In this work, beneficial mutations from random mutagenesis were mostly outfacing positions and close to or in the loops (R122I/H169Y/G49D for dCyRFP4 and V95M/Y211F for mCyRFP4), which have little direct effect on the chromophore but can tighten the packing of β-barrel to potentially enhance the rigidity of the chromophore and reduce the chance of nonradiative relaxation. The only inner facing mutation in our random mutagenesis is A104V, which was identified in both dCyRFP4 and mCyRPF4. Interestingly, A104V is also one of the three mutations that contributes to the improved brightness of mNeptune2^14^. Random mutagenesis of monomeric fluorescent proteins often results in increased homodimerization affinity, but in our case dimerizing mutants of mCyRFP4 were acceptable for use within Antares. Specifically, dCyRFP4 was generated by the mutation R122I at the dimeric interface, replacing a potentially repulsive electrostatic effects of Arg with the hydrophobic Ile. The reason for the higher expression level of dCyRFP4 and dCyRFP4-containing Antares3 compared to their predecessors may then result from faster folding and/or improved energetic stability of the dimer.

We and others^12^ have demonstrated that C166/E167 close to the C-terminus are important in tuning the catalytic activity of NLuc. Despite the higher catalytic rate against FFz we have achieved, it is challenging to further optimize the molecular brightness with existing colony-based screening given the limited throughput (10,000 colonies per day). High-throughput screening methods (e.g. colony-picker robots or single cell screening) or computer-aided deisgn and *in silico* screening may be required to examine larger libraries of variants to accelerate the evolution of NLuc-derived reporters.

Future work can expand the linkage of bioluminescent calcium sensing and biological outputs in animal brains in exciting ways. Improving RET to reducing blue emission from CaMBI may be able to expand the PhAST repertoire to include multiple color-matched luciferase/rhodopsin pairs, ultimately enabling combinatorial control of distinct neuronal populations or circuits using cell-intrinsic photon sources. As PhAST components, and the photon second messenger itself, are completely species-agnostic, the method should also be readily portable to mammalian systems. In this work, we used hikarazine, which converts slowly to furimazine, for consistency with our previous work. However, CFz or the recently optimized CFz9^8^, brain-permeant furimazine analogs that can be slowly released from poloxamer-307 hydrogels, should also work well for PhAST. Finally, a large variety of optogenetic actuators, such as light-activated transcription factors^30,31^, recombinases^32,33^, or proteases^34–36^, can be coupled downstream of bioluminescent calcium indicators, allowing for long-lived reporting, genetic manipulation, or programmable biological responses to neuronal activity, within the same neurons or nearby to them, with temporal control provided by substrate administration.

## Methods

### Cloning, mutagenesis and screening of libraries

Phusion Flash High-Fidelity PCR Master Mix (Thermo) was used for standard and overlap PCR reactions. PCR products were purified using agarose gel electrophoresis and cloned into vectors using In-Fusion HD Cloning kit (Takara Bio). Assembled constructs were transformed into XL10-Gold *E. coli* competent cells (Agilent). The cells were transferred onto an LB agar plate containing 100μg/mL Ampicilin and incubate at 37 °C overnight. To purify plasmids, single colonies were inoculated into 4 mL liquid LB medium and cultured at 37 °C for overnight. Cells were collected after centrifugation and plasmids were purified using Zyppy Plasmid Miniprep Kit (Zymo Research). Purified constructs were sequenced for verification. For site-directed mutagenesis, primers with mutations were designed for overlap extension PCR. Degenerate codon NNK or NNS was used for saturation mutagenesis at specific positions. Error-prone PCR was accomplished using GeneMorph II Random Mutagenesis Kit (Agilent). Libraries of mCyRFP or Antares were transformed into *E. coli* competent cells and cells were cultured on agar plate for protein expression and screening. mCyRFP libraries were cloned into pNCS (Allele Biotech) vectors for expression in bacteria. Colonies expressing mCyRFP variants were excited and screened using a blue light transilluminator (Vernier). Colonies exhibiting brighter or red-shifted fluorescence were subcultured in liquid LB medium. Cultured bacteria were lysed with B-PER reagent (Thermo Scientific) for scanning the fluorescence spectra in Safire2 microplate reader (Tecan).

The gene of Antares was cloned into pBAD vector for bacterial expression. 0.1% Arabinose was added into LB agar plates or liquid LB medium to induce protein expression in bacteria. To screen the bioluminescence of Antares variants in bacterial colonies, substrate FFz was sprayed onto the colonies and the plate was immediately placed into FluorChem Q with open filter and no excitation light for acquisition of bioluminescence image. Brightest colonies were picked and subcultured for further validation in bacterial lysates. Bacterial lysates were diluted 1000 times in PBS, 1:1 mixed with 50 μM FFz or CFz for bioluminescence measurement in Varioskan plate reader. FFz and CFz were generous gifts from Promega Corporation. Fluorescence was also tested in undiluted lysates and used to normalize the expression level to compare molecular bioluminescence intensity between different variants.

### Characterization of purified mCyRFPs

mCyRFP genes in pNCS vector containing His tag were transformed into *E. coli* that were transferred onto LB agar plates for overnight culture at 37 °C. Single colonies were inoculated into 100 mL LB medium for amplified protein expression. Cells were harvested by centrifugation and lysed with B-PER reagent. mCyRFP proteins were purified using Hispur Ni-NTA beads (Thermo Scientific), followed by buffer exchange to PBS using ultrafiltration tubes (Merck Millipore). Absorbance and fluorescence spectra were measured with Safire2 microplate reader (Tecan). Extinction coefficient was determined using the base-denaturation method. Quantum yield was measured using CyOFP as a standard (QY = 0.76).

### Mammalian cell culture

HEK293A cells, HeLa cells and Huh-7 cells were maintained in a humidified 5% CO_2_ incubator at 37 °C. Cells were cultured in high-glucose Dulbecco’s Modified Eagle Medium (DMEM, Life Technologies) with 5% fetal bovine serum (FBS, Life Technologies) and 2 mM glutamine (Sigma-Aldrich).

### Characterization of bioluminescent proteins in mammalian cells

Genes that encode Antares variants or mScarlet-P-GeNL were cloned into pcDNA for protein expression in mammalian cells. DMEM with or without phenol red was used for cell culture. Cells were seeded in wells of 24-well plates such that the confluency reached ~70% on the next day for transient transfection. Lipofectamine 3000 (Thermo Fisher Scientific) was used to transfect the 500 ng plasmids into mammalian cells per well. Bioluminescence was measured 24-48 h after transfection. 5× universal lysis buffer (Nanolight Technology) was diluted in PBS for cell lysis. The lysates were further diluted 100 times in PBS and 1:1 mixed with 50 μM FFz or CFz for bioluminescence measurement in Varioskan plate reader. To measure bioluminescence of luciferase in live mammalian cells, the transfected cells were detached from the well bottom with Tripsin-EDTA solution and resuspended in 300 μL DMEM (no phenol red) with 5% FBS. Cells were diluted 100 times in PBS and 1:1 mixed with 50 μM FFz for bioluminescence measurement.

### Directed Evolution of CaMBI3

The libraries of CaMBI were generated using an error prone PCR reaction using the GeneMorph II Random Mutagenesis Kit (Agilent). PCR products were purified via agarose gel electrophoresis and then cloned into the pBAD vectors using the In-Fusion Cloning Kit (Takara). The assembled constructs were then transformed into electrocompetent *E. coli* cells to express the library, which were plated on LB-agar plates supplemented with 400 μg/mL ampicillin and 0.02% (wt/vol) arabinose for overnight culture at 37 °C. On the following day, colonies were randomly selected for subsequent inoculation in 96-well plates with liquid LB medium containing ampicillin and arabinose for 18 h in a 37 °C shaking incubator. The cells were collected by centrifugation and lysed using B-PER (Thermo Scientific). The supernatant of each well was diluted into 1× Tris buffered saline (TBS), with 10 mM EGTA (0 μM calcium) or 10 mM EGTA-Ca (39 μM calcium). Luciferase substrate CFz was added to reach a final concentration of 25 μM for the luminescence measurement of the CaMBI variant in a microplate reader. The luminescence signals of each variant in 0 or 39 μM calcium were used for calculation, and plasmids of the variants exhibiting the largest signal contrast were recovered for sequencing and were used for the next round of mutagenesis.

### In Vitro Characterization of CaMBI3

CaMBI variants were cloned into pcDNA vectors for expression in mammalian cells. Assembled constructs were transfected into HEK293A cells using Lipofectamine 3000. Cells were seeded in a 24-well plate and maintained in high-glucose DMEM with 5% FBS at 37 °C in a humidified CO_2_ incubator. Then, 24 h after transfection, the medium was removed by vacuum and the cells were lysed with Passive Lysis Buffer (Promega). The cell lysates were diluted into TBS buffer with 0, 39 μM or other concentrations of calcium and the substrate was added to the buffer for luminescence measurement in a microplate reader.

### HeLa Cell Culture and Imaging of CaMBI3

HeLa cells were maintained and transfected using the same protocol as HEK293A cells (described above). Cells expressing CaMBI variants were imaged 24 or 48 h after transfection. Prior to imaging, the cell medium was replaced with HEPES buffered HBSS with 25 μM CFz substrate. Cells were imaged using a Zeiss Axiovert 200M fluorescence microscope with a 40 × 1.2-NA C-Apochromat water-immersion objective and an ORCA-Flash4.0 V2 C11440-22CU scientific CMOS camera. The microscope was controlled using MicroManager. No light source or optical filter was used for luminescence imaging. To image the fluorescence signals of CaMBI, a metal-halide arc lamp (Exfo) was used as the excitation light source with a 545RDF10 excitation filter and fluorescence was collected in a 565ALP emission filter. Time-lapse images were recorded every 5 s with 4 × 4 binning and 2 s exposure time. Histamine was added to the imaging buffer to reach a final concentration of 10 μM, approximately 1 min after the recording began.

### Culture and Imaging of Dissociated Neurons expressing CaMBI3

Dissociated rat neurons were generated according to the protocol described previously^15^. Briefly, the neurons were plated in a 24-well plate in Neurobasal media with 10% FBS, 2 mM GlutaMAX, and B27 supplement. On days-in-vitro (DIV) 1, a half of the media was changed so that the FBS can be reduced to 1% for each well. Then, a half of the media was replaced every 3 days with fresh Neurobasal media without FBS. Neurons were transfected at 9–11 DIV using Lipofectamine 2000 (Life Technologies). The imaging buffer and microscopic setup were similar to those described in Section 2.3. Time-lapse images were recorded at 10 Hz with a 0.1 s exposure time. HHBSS buffer with 50 μM KCl was 1:1 added to the cell imaging buffer to reach a final concentration of 25 μM KCl to depolarize neurons during the imaging.

### ASAP4-R4 whole-cell patch clamp recording and dual-view voltage imaging

HEK293A cells were transfected with Lipofectamine 3000 and corresponding plasmid DNAs packaged in a pCaggs backbone vector following previous protocol^37^. Briefly, 400 ng of plasmid DNA, 1 µL of P3000, and 1 µL of Lipofectamine 3000 reagents were pre-mixed in 50 µL of Opti-MEM media and incubated for 10min in room-temperature. Then the mixture was added into a well with 80% confluent HEK293A cells pre-seeded the day before. Three to four hours after the transfection, the cells were detached by adding 50 µL of 0.25% Trypsin-EDTA (Thermo Fisher Scientific) and resuspended by pipetting. Then the dissociated transfected HEK293A cells were re-plated on 12-mm diameter glass coverslips (Carolina Biological) and incubated overnight for a total of 20‒24 hr until subsequent patch clamp experiments.

Patch clamp experiment was conducted as previously described^37^. Briefly, the 12-mm diameter coverslip was placed in a patch chamber perfused with extracellular bath solution (110 mM NaCl, 26 mM sucrose, 23 mM glucose, 5 mM HEPES-Na, 5 mM KCl, 2.5 mM CaCl_2_, 1.3 mM MgSO_4_, and pH was adjusted to 7.4) in room-temperature. The glass capillary pipette pulled to have 3-5 Mohm pipette resistance was filled with intracellular solution (115 mM potassium gluconate, 10 mM HEPES-Na, 10 mM EGTA, 10 mM glucose, 8 mM KCl, 5 mM MgCl_2_ and 1 mM CaCl_2_, and pH was adjusted to 7.2). After a successful giga-ohm seal, the cell was voltage clamped using a Multiclamp 700B amplifier controlled by pClamp software while the signal was digitized by Digidata 1440B (all from Molecular device). A pulse protocol with 13 square voltage steps reaching –200, –160, –140, –120, –100, –80, –70, –60, –40, –20, 0, 30, 50, and 80 mV was applied as command voltage. Each pulse step was 400 msec long.

For excitation of the transfected cells, SOLIS-470C (Thorlabs) was used but filtered by 482/18nm bandpass filter (Semrock) and reflected by a 488nm dichroic mirror (Semrock) before the light was fed into 40× 1.3NA oil immersion lens (Zeiss). For simultaneous imaging during the whole-cell voltage clamp recording, an image splitter (W-view, Hamamatsu) equipped with a dichroic mirror (552nm) and two bandpass emission filters (525/50nm for green and 665/65nm for red) was placed before an sCMOS camera (Flash4.0 V2, Hamamatsu). The two emission channels were arranged side-by-side, so that they can be imaged on the CMOS sensor together. The sCMOS camera was operated in a 4x4 binning mode at 512×256 pixels to achieve 200 fps frame rate.

### Spectra measurements in HEK293-Kir2.1 cells

HEK293-Kir2.1 cells were maintained and transfected as previously described^37^ with following modifications. Two days before the spectra measurement, HEK293-Kir2.1 cells were plated in a 24-well glass bottom plate. Three wells were prepared for each sensor variant. In each well, 180,000 cells were seeded. After a half-day of incubation, the cells in each well were transfected with corresponding plasmid DNA in the pCaggs backbone vector by using Lipofectamine 3000. The cells were incubated overnight, and then the media in each well was replaced with the fresh culture media and incubated until the measurement. Two days after the transfection and right before the spectra measurement, the culture media was replaced with 100 µL of 10 mM HEPES supplemented HBSS for each well. The cells were detached from the glass and dissociated gently by pipetting. Subsequently, the 100 µL cell mixture was transferred to a well in a 96-well glass-bottom plate for measurements.

The spectra measurement was conducted using the Varioskan microplate reader. Excitation spectra were measured for 300‒520 nm range while measuring emission at 545 nm. Emission spectra were measured for 490‒ 700 nm range while exciting at 470 nm. There was 2 nm increment for both excitation and emission measurements. The results of three replicates (3 wells in a 96-well plate) were averaged for each sensor variant. Then the mean trace was normalized to the maximum signal before they were smoothed by 5-point running average.

### C. elegans methods

#### Growth and maintenance

*C. elegans* worms were cultivated and maintained according to standard conditions^38,39^. In short, worms were grown on regular NGM supplemented with OP50 *E. coli* strain.

#### Molecular biology

DNA for CaMBI3 was codon-optimized using the *C. elegans* codon adapter from the Max Plank Institute of Molecular Cell Biology and Genetics (https://worm.mpi-cbg.de/codons/cgi-bin/optimize.py) and one intron was introduced. DNA was synthesized and cloned into a basic plasmid by Twist Bioscience to be further moved into destination vectors using Gibson Assembly. In order to compare CaMBI1-300 and CaMBI3-970, a change in the linker region between CaM and M13 and previously described as involved in modulating calcium affinity^2^ was performed on CaMBI3-350 to make it equal to the one in CaMBI1-300 (from GGGS to G). Primers used for this directed mutagenesis are described in **Supplementary Table 2**. The *myo-3* and *rab-3* promoters for expression in BWM and neurons respectively have been described elsewhere.^40^ The constructs used for PhAST used the promoters described before.^9^ All constructs were terminated with the *let-858* 3’ untranslated region UTR originally obtained from pHW393.^41^

#### Transgenesis

Transgenic animals were generated by microinjection according to standard protocols^42^ at 10 ng/μL for plasmids either with *myo-3* or *rab-3* promoters. DNA ladder (1 kb Plus DNA Ladder, Invitrogen) to a maximum DNA load of the mix of 100–150 ng/μL was used as carrier. The integration of the resulting extrachromosomal arrays was performed as described in Das et al^43^. For the PhAST strains, 30 ng/μL of plasmid containing either CaMBI1-300 or CAMBI3-970 under the *sra-6* promoter, was used in combination with 70 ng/μL of pNP259 (*gpa-14*p::CRE, Schmitt et al, 2012) and 10 ng/μL of pCFJ68^40^ as co-injection marker. These mix was injected into the MSB985 strain (see **Supplementary Table 1**) *npr-9*p:: ChRmine::YFP::SL2::jRGECO1a was injected at 20 ng/μL with up to 100 ng/μL of DNA ladder.

#### qPCR

Sample preparation was done as described in Malaiwong et al^44^. Briefly, worms were washed off culture plates with M9 buffer, excess bacteria eliminated by successive washes and lysed in 500 µL lysis buffer supplemented with proteinase. The genomic DNA was purified using the Zymoclean Gel DNA Recovery Kit (Zymo Research) and qPCR performed using the Roche LightCycler 480 device and SYBRGreen Master mix. Primers for CaMBI1/CaMBI3 and internal control are specified in **Supplementary Table 2**.

#### Imaging

##### Luminescence

Bioluminescence imaging was performed in a home-built Low-Light microscope as described.^19^ For immobilized animals, Hikarazine Z108^45^ (Synthelis) was used as substrate. Shortly, a 2 mg/mL stock solution (40% DMSO, 60% acidic EtOH) was diluted in a buffer containing 0.2 M citric acid, 0.4 M sodium phosphate dibasic, 2 % dimethylsulfoxide (DMSO) and 0.1% Triton X-100 final concentration of 1 mg/ml. 1 µl of the mix was delivered onto the pad, worms transferred and 1 μL more added before closing the preparation with a coverslip. For imaging of moving animals, worms were transferred to a drop of buffer containing 0.2 M citric acid, 0.4 M sodium phosphate dibasic, 2 % dimethylsulfoxide (DMSO) and 0.1% Triton X-100 placed either onto 2% agarose pads (*myo-3*p::CaMBI3) or 6% agarose pads (*rab-3*p::CaMBI3). FFz 20 mM was diluted in PBS to final 1:1 proportion was used as cofactor.

#### Fluorescence

Fluorescence images were taken in the same home-built Low-Light microscope using an inverted confocal microscope (Nikon Ti2 Eclipse). Animals were imaged live in 3 mM levamisole on 2% agarose pads, and, unless otherwise specified, using a 40x/1.15 water immersion objective. The different fluorophores were imaged as follows: BFP was exited using 405 nm laser and transmitted through a 445 nm emission filter, CyOFP was excited using a 488 nm laser and transmitted through a 594 nm emission filter, YFP was exited using 514 nm laser and transmitted through a 552 nm emission filter, jRGECO1a was excited using a 561 laser and transmitted through a 594 nm emission filter. Exposure time was variable between 200–500 ms.

For fluorescence comparison of CaMBI1-300 and CaMBI3-970, worms were imaged using the same home-built Low-Light microscope used for luminescence imaging. Animals were imaged live in 3 mM levamisole on 2% agarose pads, and an LED light source emitting light at 490 nm (M490L4, Thorlabs) and ~16 µW power at the sample plane. A 600-40 nm soft-coated Bandpass filter from Thorlabs was used as emission filter. In all cases, 1 second was used as exposure time.

### PhAST

#### Plate preparation

Plates for nose-touch assays were prepared as described before^9^. Plates containing ATR were prepared by spreading 100 μL of OP50 culture mixed with ATR (0.1 mM final concentration) onto 55-mm plates containing 10 mL NGM. To prepare the plates containing Hikarazine Z108 (Synthelis), PBS needed to be prewarmed at 50 ^o^C and the 20 mg/mL stock thawed in those conditions too. 6 μL of the substrate stock were added to 200 μL of PBS, split between the control plate (no ATR) and the ATR containing plate (prewarmed at 37 ^o^C) and spread with a seeding loop until complete absortion by the agar. Plates were allowed to reach room temperature before transferring the worms.

#### Nose-touch assays

Worms were transferred to the assay plates as L4 and scored the next day as young adults. For the assays, an eyebrow hair picker placed in front of the worm for it to collide was used. A positive event was counted when, upon the contact of the tip of the nose with the hair, the worm reacted by moving backward. At least 10 worms were assayed for each condition and three experimental replicas were performed, leading to a minimum of 30 worms assayed per condition. Plates were blinded before scoring.

#### Rescue with photons

A modified version of the protocol described in Porta-de-la-Riva et al^9^ was used. Briefly, synchronized L4 animals were cultivated in the dark at 20 °C on NGM and OP50 bacteria with or without ATR. The next day worms were transferred for three hours onto plates containing either the vehicle solution for Hikarazine Z108 or the substrate itself (maintaining the ATR conditions and giving rise to 4 final conditions). Worms were transferred to assay plates after three hours and allowed to acclimatize for 10 minutes prior to the assay.

### Analysis

#### Bioluminescence intensity and dynamic range

Histograms of the luminescence intensity of animals expressing CaMBI1-300 and CaMBI3-970 were analysed in R by collecting all pixel values, corresponding to the number of photons-per-pixel recorded, between 10 and 199 counts. Pixels with 200 counts were removed to avoid undercounting saturated pixels. Fluorescence and bioluminescence images of CaMBI1-300 and CaMBI3-970 samples were processed using custom scripts in R (version 4.2.2) with the *ijtiff, ggplot2*, and *dplyr* libraries. Raw.tif image stacks were loaded from specified directories and flattened to extract single-plane intensity values. Pixel intensities from all images were pooled to define a common dynamic range, and histogram bins (n = 50) were calculated based on this global intensity distribution.

For each image, normalized pixel intensity histograms were computed by excluding saturated values and low-signal background (intensities <10 or exactly 200). These histograms were then averaged within each group (CaMBI1 or CaMBI3) to obtain group-wise distributions of pixel intensities. Density differences between the two conditions were computed for each histogram bin, and visualized both as overlaid line plots and as difference plots with LOESS smoothing. All plots were generated using *ggplot2* to compare the overall intensity distribution profiles between the two sensor constructs.

#### Curvature analysis and intensity correlation

To quantify geometric features along biological contours, we computed the local curvature of 2D shapes extracted from image data. Discrete contour coordinates (*x, y*) were first converted into arc-length parameterization, corrected by pixel scale (0.46 µm/pixel). Smoothing splines were fit to *x(s)* and *y(s)* as a function of arc length *s*, using a smoothing parameter of 0.6. First and second derivatives of the splines were then numerically evaluated to compute the curvature *κ(s)* using the standard expression:

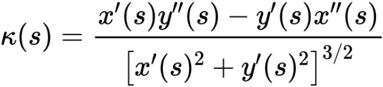

In parallel, grayscale intensity profiles were extracted along corresponding contours from calibrated microscopy images. Intensity data were smoothed using a centered rolling average (window size: 40 pixels) to reduce noise while preserving spatial trends. To compare curvature and intensity, the smoothed intensity values were interpolated onto the same arc-length grid used for curvature. We then plotted intensity as a function of curvature and visually inspected correlations between local bending and fluorescence signal. All computations were performed in R using custom scripts, with plotting and interpolation performed via base graphics and the *zoo* package.

#### Touch response contingency analysis and statistical analysis

To statistically assess differences in touch response behavior across experimental groups, binary outcome data (response: yes/no) were compiled into contingency tables comparing pairs of conditions. The comparisons included: (i) control vs. treated animals, (ii) untreated vs. mutant animals, and (iii) vehicle vs. rescue groups. For each comparison, the number of positive and negative responders was calculated using binary classification vectors, and two-by-two contingency tables were constructed.

Fisher’s exact test was applied to each table to evaluate the statistical significance of differences in response rates, particularly suitable given the small sample sizes and categorical nature of the data.

To visualize these results, all contingency tables were combined and a balloon plot was generated in R, with point size representing the count in each cell and numeric labels overlaid for clarity. This visualization facilitated comparison across groups and conditions by displaying relative frequencies of responses in a compact and intuitive format.

For each animal, touch responsiveness was quantified as the sum of positive responses across ten repeated trials. These cumulative scores were computed for multiple experimental groups (e.g., treated, vehicle, wild-type, mutants) and organized into a unified data frame for statistical comparison. The resulting dataset was visualized using jittered scatter plots and summary statistics (mean ± 95% CI).

To assess differences in median response counts, nonparametric bootstrapping (resampling with replacement) was employed. Median differences between control and test groups were calculated over 1,000 iterations, and confidence intervals were derived using both exact and bootstrap-based methods. A density plot was used to visualize the distribution of resampled median differences.

## Author contributions

Y.Z., M.P.R., M.K. (Krieg), and M.Z.L. designed experiments. Y.Z. engineered and characterized CyRFP4 variants, Antares3 and CaMBI3 variants. M.P.R. characterized CaMBI3 in *C. elegans* and performed the experiments in the nematode. S.L. and Y.W. characterized ASAP4-R4, and CyRFP4, respectively. Y.Z., M.K. (Klein), M.P.H., and

Y.S. characterized Antares3. Y.Z., M.P.R., S.L., Y.W., M.P.H. analyzed the data. L.P.E., T.A.K., M. K. (Krieg), and M.Z.L. provided supervision. Y.Z., M.P.R., M. K. (Krieg) and M.Z.L. edited the manuscript.

## Acknowledgements

The work was supported by a Stanford School of Medicine Dean’s Fellowship (Y.Z.), a Stanford Discovery Innovation Award (M.Z.L.), NIH grants 1R01NS123681, R21NS122055, R21DA048252 (M.Z.L.), CNS2022-135906 funded by MCIN/AEI/ 10.13039/501100011033 and by the “European Union NextGenerationEU/PRTR (M.K.) and European Union (ERC PoC, 101138041) as well as “Severo Ochoa” program for Centres of Excellence in R&D (CEX2019-000910-S), the Fundació Privada Cellex, Fundació Mir-Puig, and from Generalitat de Catalunya through the CERCA and Research program (M.K.). We thank Prof. Robert E Campbell at the University of Tokyo for his support and feedback during the experimental work of CaMBI3 evolution.

