## Supplementary Information for "Bright calcium-modulated bioluminescent indicators for activity imaging and red photon-assisted synaptic transmission"

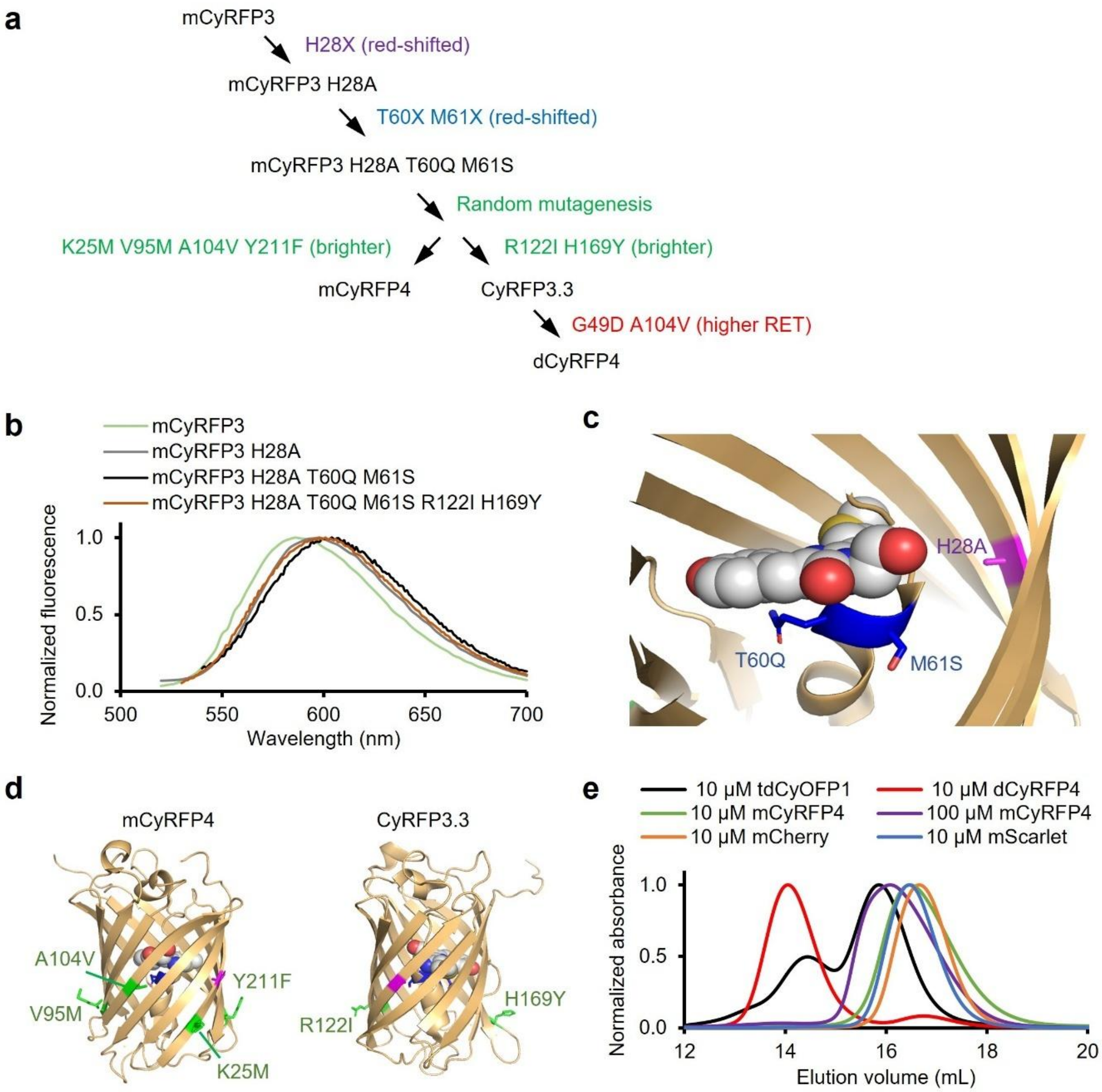


**Supplementary Figure 1. Directed evolution of mCyRFP3. a**, Evolution trajectory of dCyRFP4 and mCyRFP4. **b,** Emission spectra of mCyRFP3 variants with bathochromic shift. **c**, Model of the red-shifted mCyRFP3 H28A T60Q M61S based on the crystal structure of CyOFP (PDB ID 5BQL). **d**, Mutations found to improve brightness or expression in mCyRFP4 and CyRFP3.3. **e**, Size exclusion chromatography of purified mCyRFP4, dCyRFP4 and other fluorescent proteins.

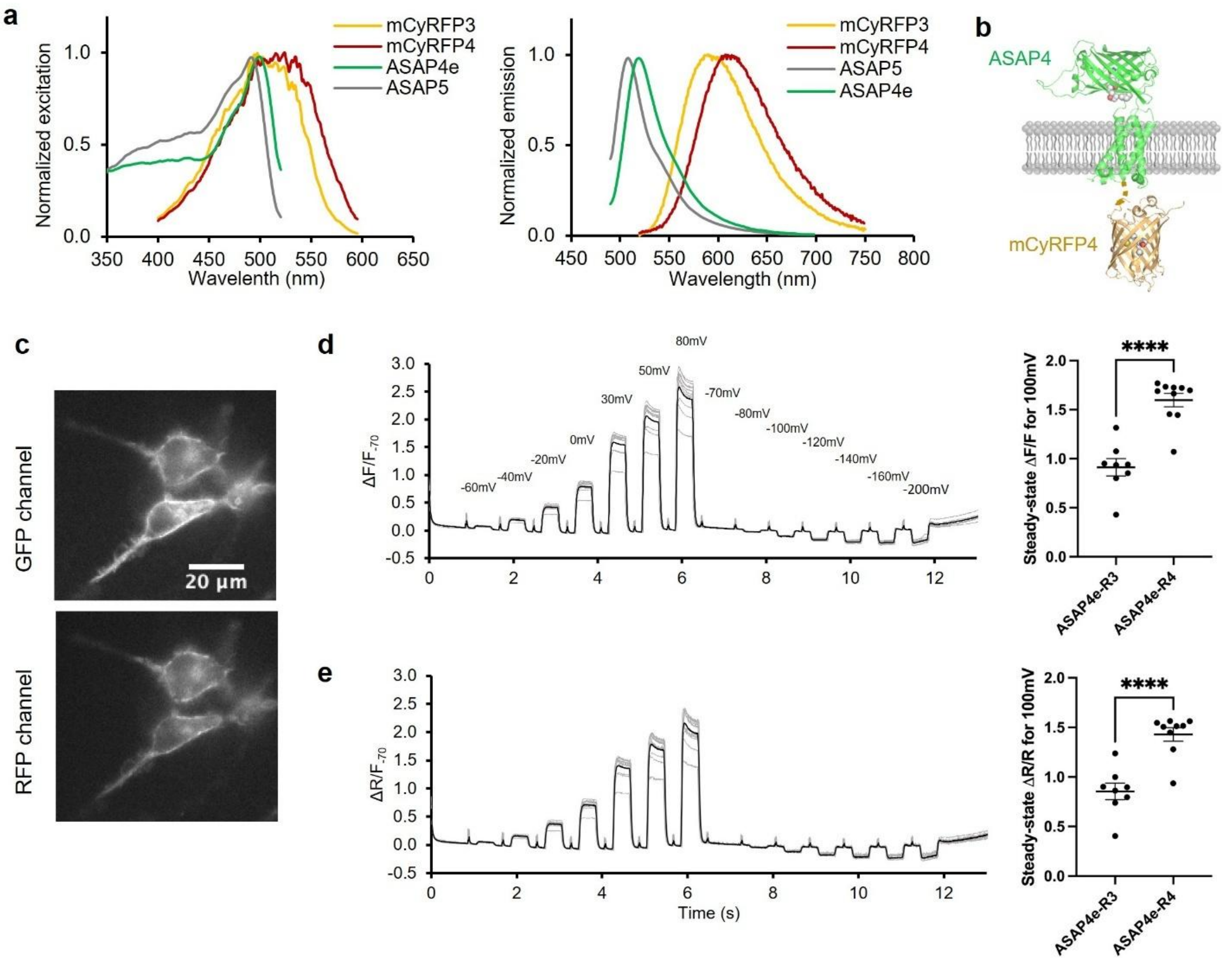


**Supplementary Figure 2. A ratiometric positively tuned voltage indicator, ASAP4e-mCyRFP4 (ASAP4e-R4). a,** Excitation and emission spectra of ASAP4e, ASAP5, mCyRFP3 and mCyRFP4. **b,** Schematic structure of ASAP4-mCyRFP4. **c,** Representative fluorescence microscopic images of ASAP4e-mCyRFP4 expressed in HEK293A cells. **d,** Left, Steady-state intensiometric responses of ASAP4e-R4 in GFP channel to voltage in HEK293A cells. Signals are normalized to -70 mV. Traces from individual cells in light grey (n=10), and the average trace in black. Right, ΔF/F of ASAP4e-R3 and ASAP4e-R4 in response to 100 mV change (-70 mV to 30 mV) in GFP channel. **e**, Left, steady-state red/green ratiometric responses (dual emission model) of ASAP4e-R4 to voltage in HEK293A cells. Signals are normalized to -70 mV. Right, ΔR/R of ASAP4e-R3 and ASAP4e-R4 in response to 100 mV change (-70 mV to 30 mV). **** $p < 0.0001$ unpaired t-test.

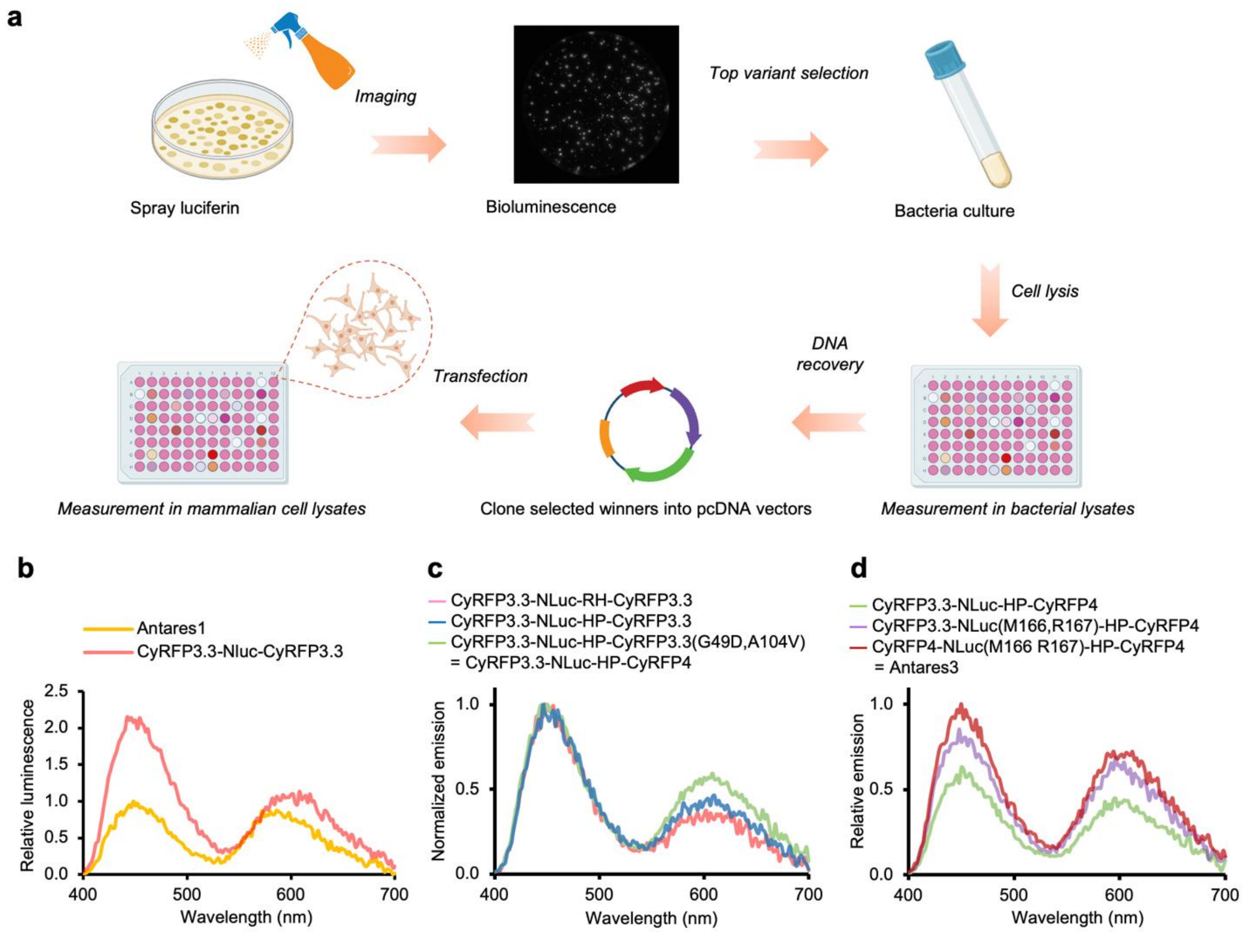


**Supplementary Figure 3**. **Optimization of Antares1 reporter. a**, Workflow for screening libraries of linkers constructed through NNK degenerate codons or libraries generated through random mutagenesis. **b–d,** Luminescence spectrum of intermediates between Antares1 and Antares3.

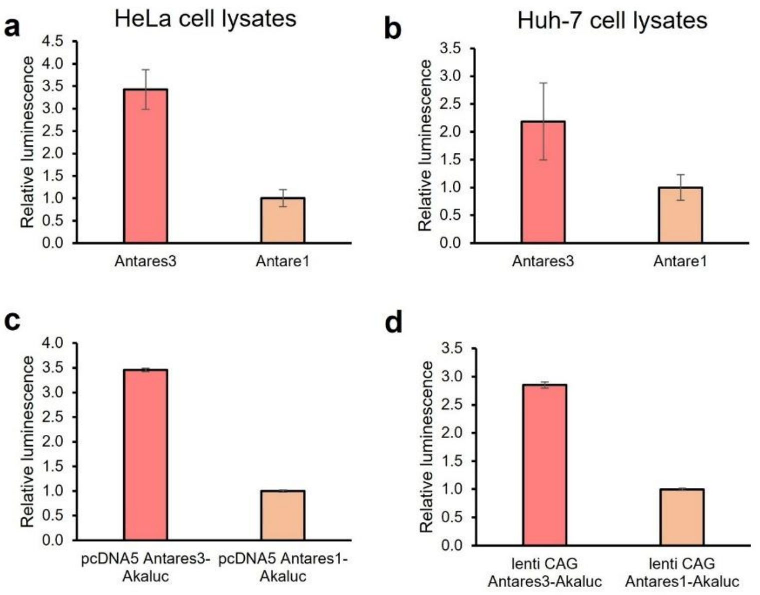


**Supplementary Figure 4. Comparison of Antares1 and Antares3 in different mammalian cell lines and in different vectors. a-b**, Luminescence collected in the diluted lysates of HeLa cells (a) and Huh-7 cells (b). Cells were transfected with the same amount of DNA. pcDNA vector was used. **c-d**, Luminescence collected in the diluted lysates of HEK293A cells transfected with the same amount of DNA (different plasmid vectors as indicated in each panel). Luminescence was recorded at a rate of once per minute for 10 minutes, and the integrated signals were calculated and compared. Error bars stand for standard deviation.

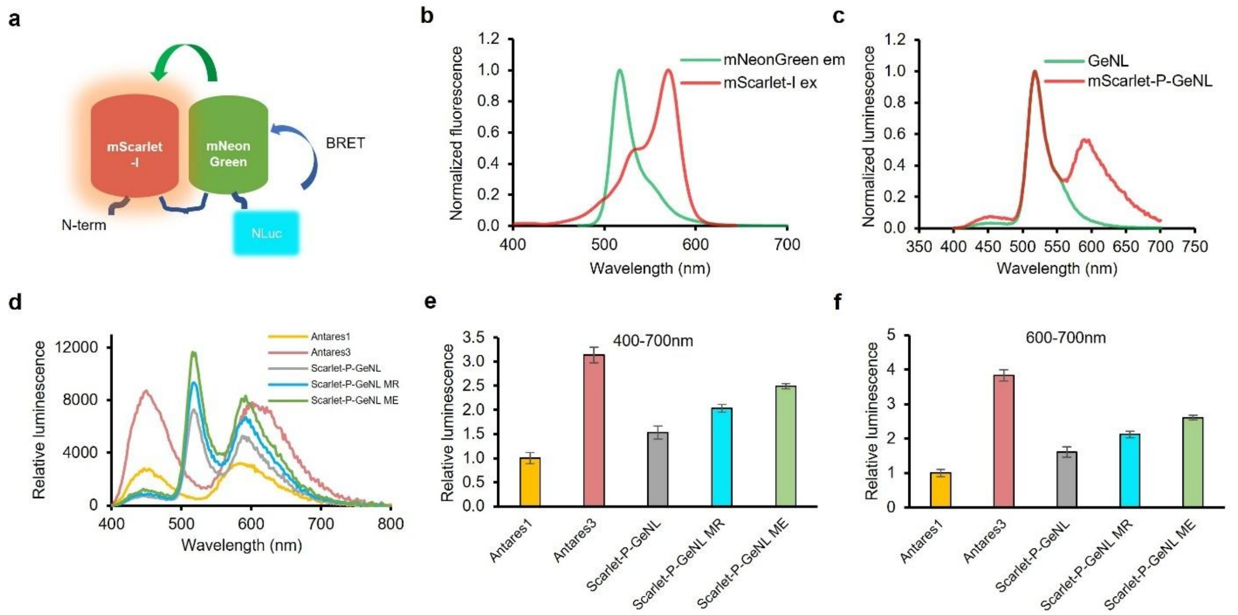


**Supplementary Figure 5. Characterization of Scarlet-P-GeNL variants. a.** Schematic structure of GeNL-P-mScarlet. **b.** Excitation spectrum of mNeonGreen and emission spectrum of mScarlet-I. Data from Fluorescent Protein Database (fpbase.org) **c.** Normalized luminescence spectra of GeNL and Scarlet-P-GeNL. **d.** Luminescence spectra of Scarlet-P-GeNL and Antares variants measured in diluted HEK293A lysates. Cells were transfected with the same amount of plasmid. **e**-**f**. Relative intensity of total and red luminescence of Scarlet-P-GeNL and Antares variants in diluted HEK293A lysates. Luminescence was measured as single time point spectrum scanning and calculated as integrated intensity from 400-700 nm (e) or 600-700 nm (f). Scarlet-P-GeNL MR represents Scarlet-P-GeNL C166M E167R, and Scarlet-P-GeNL ME represents Scarlet-P-GeNL C166M. The color of each column corresponds to that in (d). Error bars represent standard deviation.

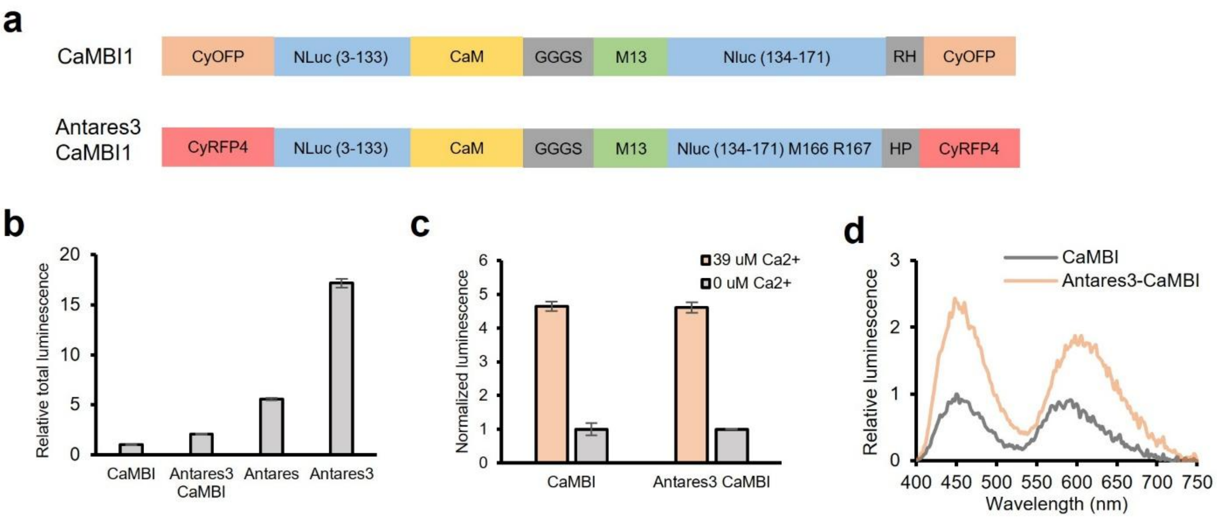


**Supplementary Figure 6. Generalization of Antares3 mutations to CaMBI using CFz. a**, Schematic domain structure of CaMBI1 and Antares3 CaMBI (CaMBI2.5). **b-d,** Relative luminescence of Antares and CaMBI variants in diluted HEK293A cell lysates in 0 or 39 µM $Ca^{2+}$ buffer. Cells were transfected with the same amount of plasmid. Total luminescence in the presence of 39 µM $Ca^{2+}$ (b) was recorded at a rate of once per minute for 10 minutes, and the integrated signals were calculated and compared. The dynamic range of the original CaMBI (Antares1 based) and Antares3 CaMBI was compared in the presence of 0 or 39 µM $Ca^{2+}$ buffer (c). Luminescence spectra (d) of CaMBI and Antares3 CaMBI were recorded at a single time point. Relative intensity indicates the signal in the diluted lysate of HEK293A cells in the presence of 39 µM $Ca^{2+}$. Error bars stand for standard deviation.

**a**

MVSKGEELIKENMRSKLYLEGSVNGHQFKCTAEGEGKPYEGKQTARIKVVEGDPLPFAF
DILAQSFMYGSKVFIKYPADLPDYFKQSFPEGFTWERVMVFEDGGVLTVTQDTSLQDGEL
IYNVKLIGVNFPANGPVMQKKTLGWEPSTETMTPADGGLEGRCDKVLKLVGGGYLHVNF
KTTYKSKKPVKMPGVHYVDRRLERIKEADNETYVEQYEHAVARYSNLGGGFTLEDFVGD
WRQTAGYNLDQVLEQGGVSSLFQNLGVSVTPIQRIVLSGENGLKIDIHVIIPYEGLSGDQM
GQIEKIFKVVYPVDDHHFKVILHYGTLVIDGVTPNMIDYFGRPYEGIAVFDGKKITVTGTLIM
HDQLTEEQIAEFKEAFSLFDKDGDGTITTKELGTVMRSLGQNPTEAELQDMINEVDADGN
GTIYFPEFLTMMARKMKDTDSEEEIREAFRVFDKDGNGYISAAQLRHVMTNLGEKLTDEE
VDEMIREADIDGDGQVNYEEFVQMMTAKGGGSKRRWKKNFIAVSAANRFKKISSSGALE
LWNGNKIIDERLINPDGSLLFRVTINGVTGWRLMRRILAHPELIKENMRSKLYLEGSVNGH
QFKCTAEGEGKPYEGKQTARIKVVEGDPLPFAFDILAQSFMYGSKVFIKYPADLPDYFKQ
SFPEGFTWERVMVFEDGGVLTVTQDTSLQDGELIYNVKLIGVNFPANGPVMQKKTLGWE
PSTETMTPADGGLEGRCDKVLKLVGGGYLHVNFKTTYKSKKPVKMPGVHYVDRRLERIK
EADNETYVEQYEHAVARYSNLGGGMDELYK

**b**

Mutations identified in screening round 1: L393I, D439E, G544D
Mutations identified in screening round 2: G329R, R515M (+previous mutations in round 1)
Mutations identified in screening round 3: N427D, V482I, A538T (+previous mutations in rounds 1 and 2)

**Supplementary Figure 7. Sequence of the CaMBI2.5 with highlighted mutations towards CaMBI3-350. a,** CyRFP4 domains (red), NLuc domain (blue), CaM domain (yellow), M13 (green), and linkers (black). **b**, Summary of mutations identified in each round of screening.

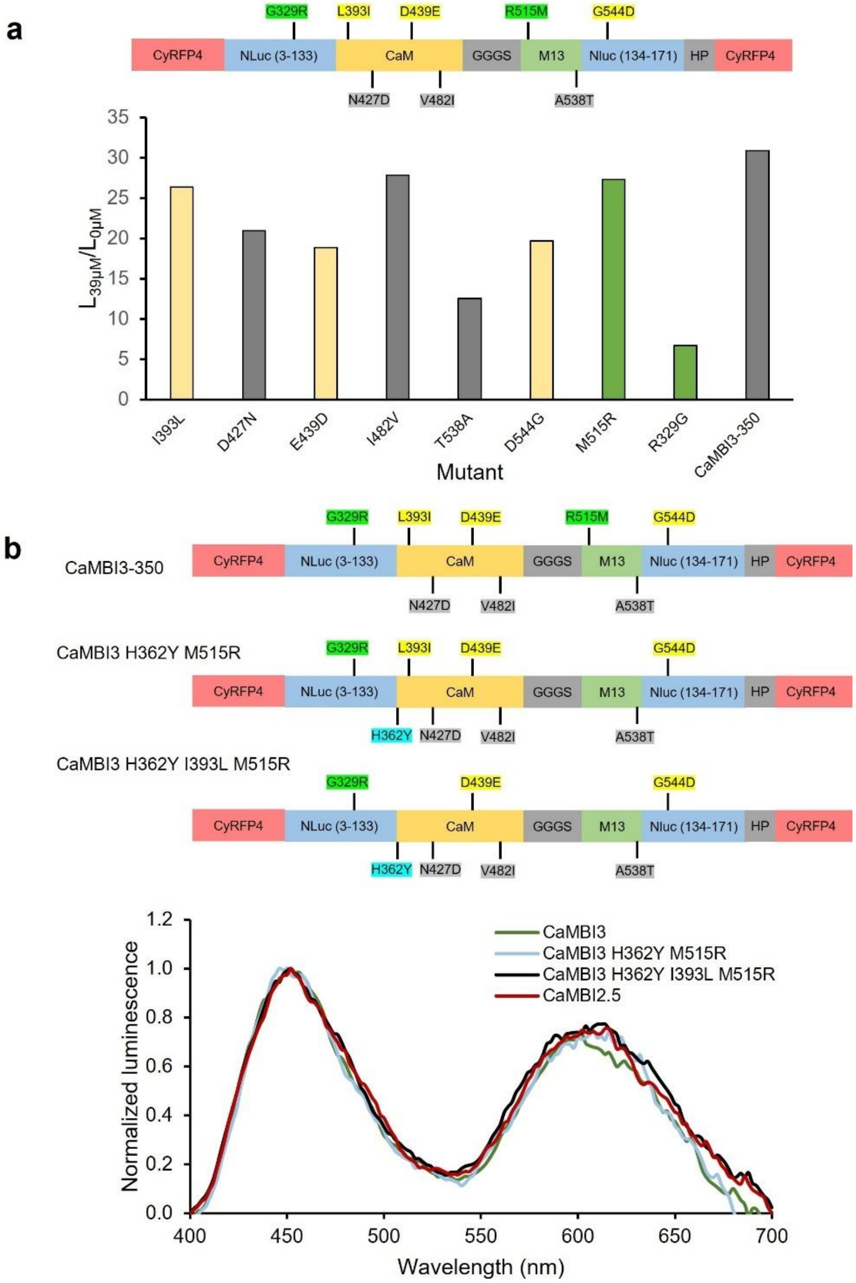


**Supplementary Figure 8. Characterization of CaMBI3-350 variants. a**, Top, Schematic domain structure of CaMBI3-350 with highlighted mutations relative to CaMBI2.5. Bottom, Reversion of single mutation in CaMBI3-350 and the dynamic range in diluted bacterial lysates in the presence and absence of $Ca^{2+}$. Luminescence was recorded at a sampling rate once per min and integrated over five time points for comparison. The color of each column corresponds to the highlighted mutations above. **b**, Top, Schematic domain structure of CaMBI3-350 variants with highlighted mutations relative to CaMBI2.5. Bottom, Luminescence spectrum of CaMBI3-350 variants measured in diluted HeLa cell lysate in the presence of $Ca^{2+}$ at a single time point.

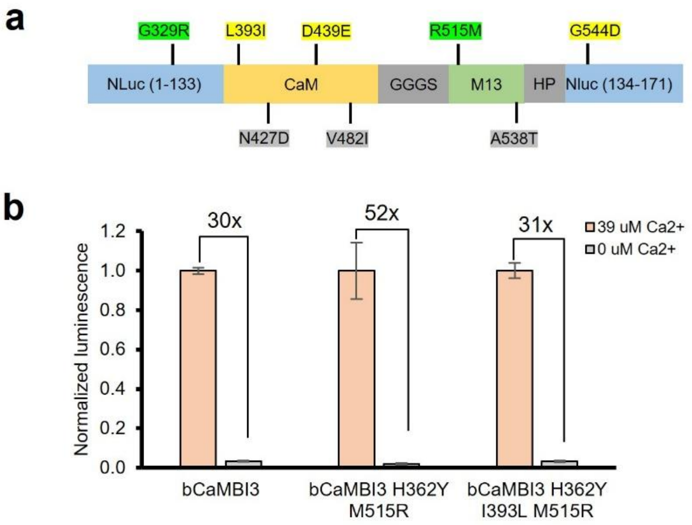


**Supplementary Figure 9. Generation of blue CaMBI3 (bCaMBI3) variants. a**, Schematic domain structure of bCaMBI3 with mutations relative to bCaMBI1. **b**, Dynamic range of bCaMBI3 variants in bacterial lysates. Luminescence was recorded at a sampling rate once per min and integrated over 5 min for comparison. Error bars represent standard deviation.

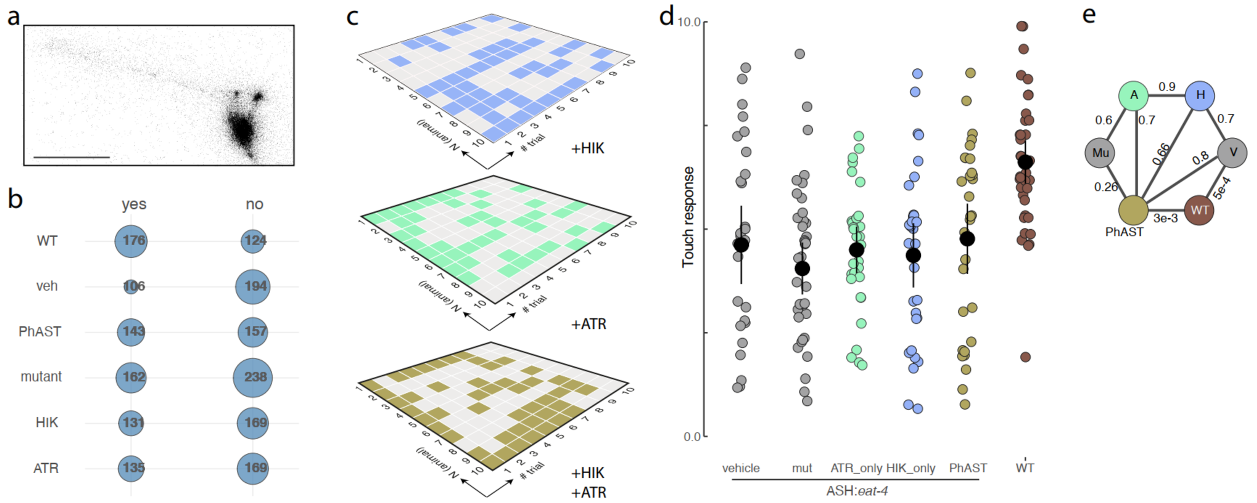


**Supplementary Figure 10.** ***CaMBI1-300 does not enable PhAST.*** **a**, Representative bioluminescent image of ASH expressing CaMBI1-300 acquired at 1s exposure time as described in the methods. Scale bar = 50 µm. **b**, Contingency matrix for all conditions tested with CaMBI1-300. Selected p-values derived from Fisher's exact test are indicated on the bracket to the right. **c**, Representative raster plots showing the distribution of responses and no responses for the three indicated conditions. Note, no history effect or habituation is observed for individual animals touched consecutively. **d**, Scatterplot showing the average nose touch responses of each animal to ten touches carrying a conditional *eat-4* KO defect in ASH and expresing CaMBI1-300 in ASH. Individual datapoints (n = 30 animals) are displayed for each condition. Each data point corresponds to the average of ten touch tests per animal. Brown data points indicate wild-type animals. Black points indicate the median; vertical bars indicate 95% CI. Floating axes indicate the bootstrapped distribution of the Paired Median Difference (PMD), the median as a red point ± 95% CI as vertical bars. **e**, Statistics between the indicated conditions in which the numbers indicate the two-sided P value derived from a pairwise comparison of the indicated conditions with a non-parametric Dunn test.

**Supplementary Table 1.** *C. elegan* strains used in this study.

| Strain name | Genotype |
|---|---|
| MSB985 | *mirSi37[flp-18p::lox2272::mtagBFP2::tbb-2 3'::lox2272::ChRmine::SL2::jRGECO1a::let-858 3']+Cbrunc-119(+)]* |
| MSB1370 | *oxTi1022[eft-3p::gfp::2xNLS::tbb-2 3' + NeoR] I; lite-1(ce314), bus-17(br2) X; mirEx566[myo-3p::CaMBI3-350::let-858 3']* |
| MSB1381 | *lite-1(ce314), bus-17(br2) X; mirEx574(rab-3p::CaMBI3-350::his-58::let-858 3')* |
| MSB1393 | *unc-119(ed3) III; mir98[*oxTi553[eft-3p::tdTomato::his-58::unc-54 3' + Cbr-unc-119(+)]] V; mirIs229[myo-3p::CaMBI1-300::let-858 3']* |
| MSB1398 | *unc-119(ed3) III; mir102[*oxTi553[eft-3p::tdTomato::his-58::unc-54 3' + Cbr-unc-119(+)]] V; mirIs234[myo-3p::CaMBI3-970::let-858 3']* |
| MSB1399 | *lite-1(ce314), bus-17(br2) X; mirIs234[myo-3p::CaMBI3-970::let-858 3']* |
| MSB1402 | *lite-1(ce314), bus-17(br2) X; mirIs234[myo-3p::CaMBI1-300::let-858 3']* |
| MSB1416 | *lite-1(ce314), bus-17(br2) X; mirIs240(rab-3p::CaMBI3-350::his-58::let-858 3')* |
| MSB1469 | *mirSi37[flp-18p::lox2272::mtagBFP2::tbb-2 3'::lox2272::ChRmine::SL2::jRGECO1a::let-858 3']+Cbrunc-119(+)], mir12[loxP eat-4], mir17[eat-4+loxP intron2] X; mirIs249[npr-9p::ChRmine::yfp::SL2::jRGECO1a+Cbrunc-119(+)]; mirEx597[sra-6p::CaMBI1-300, gpa-14p::CRE, unc-22p::gfp+Cbrunc-119(+)]* |
| MSB1470 | *mirSi37[flp-18p::lox2272::mtagBFP2::tbb-2 3'::lox2272::ChRmine::SL2::jRGECO1a::let-858 3']+Cbrunc-119(+)], mir12[loxP eat-4], mir17[eat-4+loxP intron2] X; mirIs249[npr-9p::ChRmine::yfp::SL2::jRGECO1a+Cbrunc-119(+)]* |
| MSB1481 | *mirSi37[flp-18p::lox2272::mtagBFP2::tbb-2 3'::lox2272::ChRmine::SL2::jRGECO1a::let-858 3']+Cbrunc-119(+)], mir12[loxP eat-4], mir17[eat-4+loxP intron2] X; mirIs249[npr-9p::ChRmine::yfp::SL2::jRGECO1a+Cbrunc-119(+)]; mirEx600 [sra-6p::CaMBI3-970, gpa-14p::CRE, unc-22p::gfp+Cbrunc-119(+)]* |

**Supplementary Table 2.** Primers used in *C. elegan* study.

| Target | Primers | Aim |
|---|---|---|
| CaMBI1/CaMBI3-350 | FW: ATGGTTTCTAAGGGAGAAGAACTTATCAAGG<br>RV: CTTGTAAAGCTCATCCATTCCTCCTCC | Gibson assembly |
| CaMBI3-350-GGGS linker to G | FW: TCAAATGATGACTGCTAAAGGAAAGATGCGTTGGAAGAAGAATTTCATCGC<br>RV: AATTCTTCTTCCAACGCATCTTTCCTTTAGCAGTCATCATTTGAACGAACTCC | Directed mutagenesis |
| CaMBI1/CaMBI3-350 | FW: CCAGTCGACGACCACCACTTCAAG<br>RV: GGGTTCCAGTAACGGTGATCTTCTTTCC | qPCR |
| *rps-25* | FW: TCACACCATCCGTCGTCTCTG<br>RV: GACCTGTCCGTGATGATGAACG | |
